# Tet2 Controls β cells Responses to Inflammation in Type 1 Diabetes

**DOI:** 10.1101/2020.09.01.278028

**Authors:** Jinxiu Rui, Songyan Deng, Gerald Ponath, Romy Kursawe, Nathan Lawlor, Tomokazu Sumida, Maya Levine-Ritterman, Ana Luisa Perdigoto, Michael L. Stitzel, David Pitt, Jun Lu, Kevan C. Herold

**Affiliations:** Departments of Immunobiology and Internal Medicine, Yale University, New Haven, CT 06520, USA; Department of Neurology, Yale School of Medicine, New Haven, CT, 06511, USA; The Jackson Laboratory for Genomic Medicine, Farmington, CT 06032, USA; Department of Genetics and Genome Sciences and Institute for Systems Genomics, University of Connecticut, Farmington, CT 06032, USA; Department of Genetics, Yale University, New Haven, CT 06520, USA

**Author notes:** Correspondence to: Kevan C. Herold, MD, CNH Long Professor of Immunobiology and Internal Medicine, Yale University, 300 George St, #353E, New Haven, CT 06520.

## Abstract

β cells may participate and contribute to their own demise during Type 1 diabetes (T1D). We identified a novel role of Tet2 in regulating immune killing of β cells. Tet2 is induced in murine and human β cells with inflammation but its expression is reduced in surviving β cells. Tet2-KO mice that receive WT bone marrow transplants develop insulitis but not diabetes and islet infiltrates do not eliminate β cells even though immune cells from the mice can transfer diabetes to NOD/*scid* recipients. Tet2-KO β cells show reduced expression of inflammatory genes, associated with closed transcription factor binding sites. Tet2-KO recipients are protected from transfer of disease by diabetogenic immune cells. We conclude that Tet2 regulates pathologic interactions between β cells and immune cells and controls intrinsic protective pathways. Modulating TET2 may enable survival of β cells or their replacements in the setting of pathologic immune cells.

## Introduction

Type 1 diabetes (T1D) is a chronic autoimmune condition that occurs over years after the first signs of autoimmunity which are represented by the appearance of autoantibodies. With the improvement in the sensitivity of measurement of C-peptide and the availability of tissue sections from individuals who have died with T1D, it has become clear that not all β cells are destroyed by autoimmunity. Even 50 years after the diagnosis of T1D about 2/3 of patients still have detectable levels of C-peptide indicating residual β cells. Moreover, the presence of detectable levels of proinsulin suggests dysfunctional but persistent β cells. The reasons why some β cells survive and others succumb to autoimmune killing are not certain.

There is growing evidence showing β cells are not passive bystanders of their own destruction, but instead participate and contribute to their own demise in the pathogenesis of autoimmune diabetes. Genome-wide association studies (GWAS) show that >50% of gene loci associated with T1DM are expressed in β cells. These studies, coupled with emerging molecular evidence that β cells are impaired by gain-of-function or loss-of-function of these loci, suggest an active role for the β cell in eliciting its own demise ^1–4^. The ways in which β cells participate may involve more than just their own cell intrinsic properties. Accumulating data suggests that immune tolerance to β cells reflects the physiological interaction between the immune system and β cells rather than just the properties of tolerant lymphocytes. The presence of β cells is required for initiation of diabetes autoimmunity to proceed. Spleen cells from β-cell-deprived NOD mice cannot transfer diabetes although they maintain immune competency. Furthermore, already committed “diabetogenic” spleen cells showed a reduced capacity to transfer diabetes after transient “parking” in β-cell-deprived mice ^5^. The islet cell mass plays a critical role in diabetes – surgical removal of 90% of pancreatic tissue before onset of insulitis induced a long-term protecion in NOD mice.

Lymphocytes from pancreatectomized diabetes-free mice exhibited low response to islet cells ^6^. Curiously, in NOD mice and human type 1 diabetes, autoimmunity does not extend to cells that share antigens with β cells, particularly cohabitating endocrine islet cells and neuronal cells that express GAD and IA-2. It is thus likely that both the expression by target cells of autoantigens and their presentation under conditions that allow recognition by T cells are prerequisites to the development of autoimmune lesions. Thus, the interplay between β cells and immne cells determine the outcome of autoimmunity.

β cells are also known to be intrisincally vulnerable, and are more prone than other islet endocrine cells, to death under ER and immune stress conditions ^7 8^. Early physiological β cell death triggers priming of autoreactive T cells by dendritic cells in pancreatic lymph nodes (pLNs) and mice are protected from diabetes after preventive excision of pLNs at 3 weeks of age, providing direct evidence for the importance of the local presentation of autoantigens in the diabetes process ^9,10^. Thompson et al. described a subpopulation of β cells that become senescent and actively promotes the immune-mediated destruction process ^11^. Recently Lee et al. reported that modulating the unfolded protein response (UPR) in NOD β cells by deleting the UPR sensor IRE1α prior to insulitis induced a transient dedifferentiation of β cells, resulting in substantially reduced islet immune cell infiltration and β cell apoptosis and protection from diabetes ^12^. When we analyzed β cells during the progression of diabetes, we identified a subset of β cells with dedifferentiated features such as reduced expression of *Pdx1, Nkx6.1, MafA*, and *Ins1, Ins2* as well as β cell autoantigens and those cells were resistant to immune mediated killing ^13^.

Among the mechanisms that might account for the changes of β cells in response to stressors are epigenetic changes involving silencing or activation of genes as a consequence of signaling by factors such as inflammatory cytokines. Stefan-Lifshitz et al associated DNA hydroxy methylation by IFNα ^14^.We previously described epigenetic changes in β cells from NOD mice that led to methylation marks in *Ins1* and *Ins2* and reduced gene transcription and similar changes in *INS* when human islets were cultured with inflammatory cytokines ^15^. Accordingly, we identified increased expression of Dnmts that we postulated might result in methylation of CpG sites in the insulin genes and repression of gene transcription ^16^. In addition, the responses to inflammation and epigenetic modifications may be linked by the activity of the ten eleven translocation (Tet) methylcytosine dioxygenase family members. Zhang et al showed that Tet2 is required to resolve inflammation by recruiting Hdac2 to repress IL-6. This activity of Tet2 was independent of its modifications of the epigenome ^17^. In mice, Tet2 sustained the immunosuppressive function of tumor-infiltrating myeloid cells to promote melanoma progression ^18^. In melanoma and colon tumor cells deletion of Tet2 reduced chemokine expression and tumor infiltrating lymphocytes, enabling tumors to evade antitumor immunity and to resist anti–PD-L1 therapy ^19^. In primary microglial cells lacking Tet2, there is inhibition of IL-6 release and TNF-α after LPS treatment ^20^. Deletion of Tet2 in T cells decreased their cytokine expression, associated with reduced p300 recruitment ^21^. On the other hand, enhancing TET activity with ascorbate/vitamin C increased chemokines, tumor-infiltrating lymphocytes and anti-tumor immunity ^19,22^. Lastly, a role for Tet2 in cell survival has also been suggested as Tet2-KO hematopoietic stem and progenitor cells have reduced apoptosis ^23^. These studies suggested a strategy that might be developed to protect β cells from immune damage.

Despite the changes in β cells, the control of these adaptive responses by epigenetic mechanisms and, importantly, the relationship between the changes in β cells and the control of the immune response against them has not been known. Therefore, we analyzed epigenetic changes in β cells from mice and humans during inflammation and with autoimmunity.

## Results

### Induction of Tet2 in Murine β Cells during Progression of Autoimmune Diabetes

To understand whether and how epigenetic modifications may change β cells during autoimmune diabetes, we studied gene expression changes in whole islets and β cells from the NOD model. We first analyzed the expression of Tet genes by RT-PCR. Soon after weaning, at a period of time that has been associated with the initiation of autoimmunity, we found a modest increase in expression of *Tet1* but substantially increased expression of *Tet2* in whole islets (**Fig. 1**a). In B6 mice of the same age, we did not find a similar change in the expression of Tet genes. Our earlier studies of epigenetic changes had indicated that inflammatory cytokines could stimulate epigenetic changes even in islets from normal B6 mice and we also found that there was increased expression of *Tet1* and *Tet2* but not T*et3* in islets from B6 mice following culture with cytokines (**Fig. 1**b). To determine whether these changes in Tet genes occurred in β cells, we enriched for β cells by sorting on Zinc+ and TMRE+ islet cells and found that *Tet2* expression was increased in β cells from 8- and 11-wk-old NOD mice compared to 3-wk-old NOD (**Fig. 1**c). The changes in *Tet1* expression were modest in the same β cells. The increased *Tet2* gene transcription was associated with increased intracellular Tet2 expression in NOD β cells, analyzed by FACS, at 4 wks and at the time of hyperglycemia when there is significant insulitis. We used Tet2-KO mice as a control for staining specificity (**Fig. 1**d).

**Fig 1:**
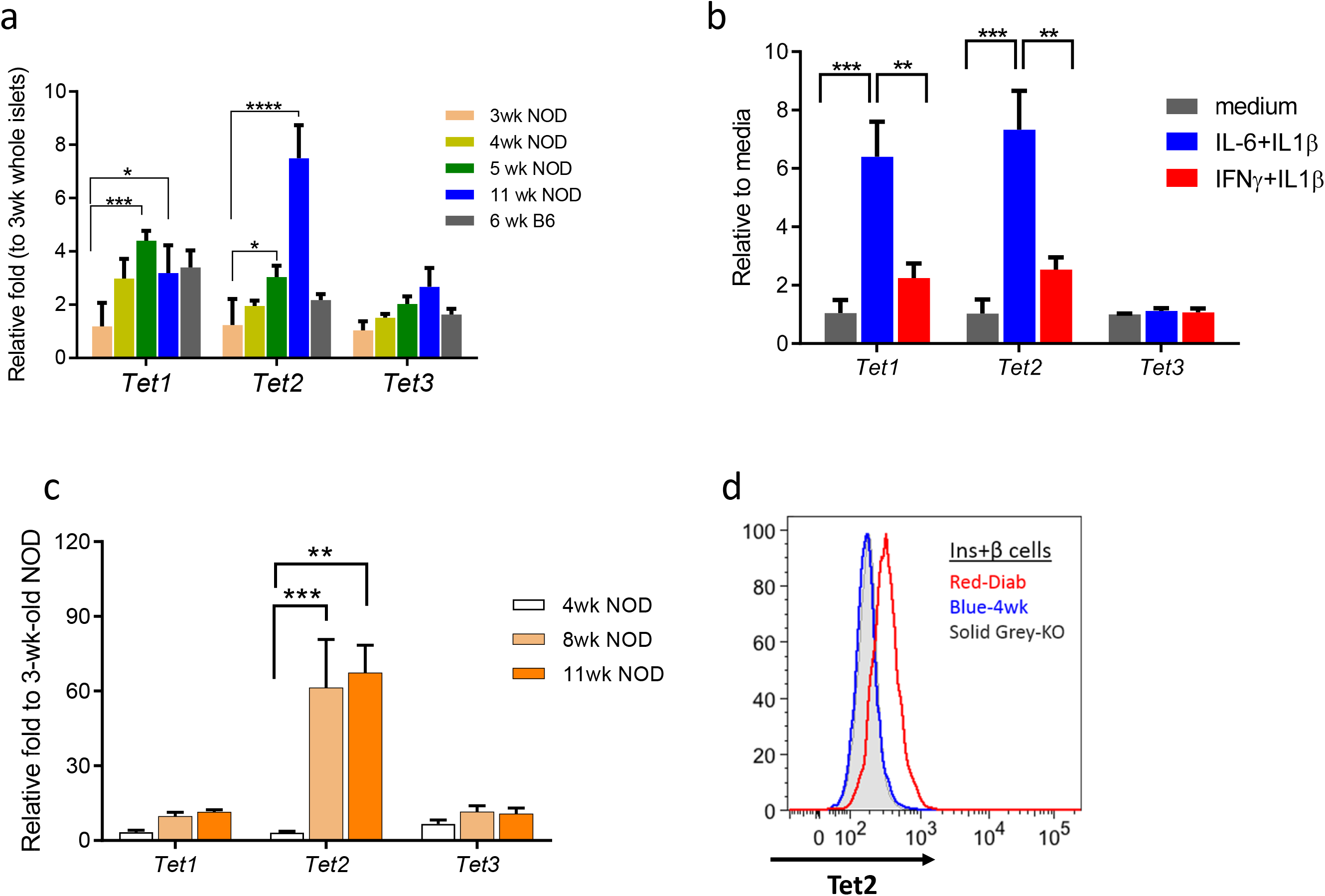
Increased Tet2 expression in islet β cells during diabetes progression in non-obese diabetic (NOD) mice. Transcription analysis of the *Tet* genes in (a) hand-picked islets from NOD and B6 mice of different age, (b) islets from B6 mice following 24-hr culture with indicated cytokines and (c) enriched β cells (Zn+TMRE+) sorted from NOD mice of different ages. RNA was recovered and *Tet* genes were measured by qRT-PCR. The Ct values were normalized to *Actb* mRNA levels (ΔC_t_=C_t_ of *Actb*-C_t_ of *target gene*+ 20). Data show the mean ± SEM of three experiments, each with 3-6 mice. T-tests were performed with FDR of 5% for multiple comparisons (\**p*<0.05, **p<0.01, \*\*\**p*<0.001, \*\*\*\**p*<0.0001). (d) Histogram comparing Tet2 protein level in β cells from 4-wk-old as well as new-onset diabetic NOD mice analyzed by FACS. β cells from NOD Tet2-KO mice were included as negative control for Tet2 staining. Tet2+ β cells were identified by intracellular staining with antibodies against insulin and Tet2. Data are representative of 3 experiments, *n*=3-5 for each experiment.

### Expression of TET2 in Human β Cells

We also studied *TET2* expression in human islets from different inflammatory settings by immunohistochemistry. Because of the profound increase in *Tet2* expression in NOD β cells we focused our studies on this epigenetic regulator. In control pancreases, TET2 expression was seen outside of the islets among exocrine tissue (**Fig. 2**a (a)). However, there was increased TET2 expression in the nuclear and cytoplasm in β cells from a patient with autoimmune pancreatitis (with cellular immune infiltrates) (**Fig. 2**a (b)), as well as a patient with T1D autoantibodies but not diagnosed with T1D (**Fig. 2**a (c)), and a patient with recent onset T1D (**Fig. 2**a (d)). The expression of TET2 was associated with inflammatory lesions since we did not identify it in the β cells in a pancreas from a patient with chronic pancreatitis in which infiltrating immune cells were not found (**Fig. 2**a (f)). Interestingly, we also did not identify TET2 staining in the remaining singular β cells in the pancreas from a patient with long-established T1D who did not have detectable autoantibodies or insulitis (**Fig. 2**a (e)).

**Fig. 2:**
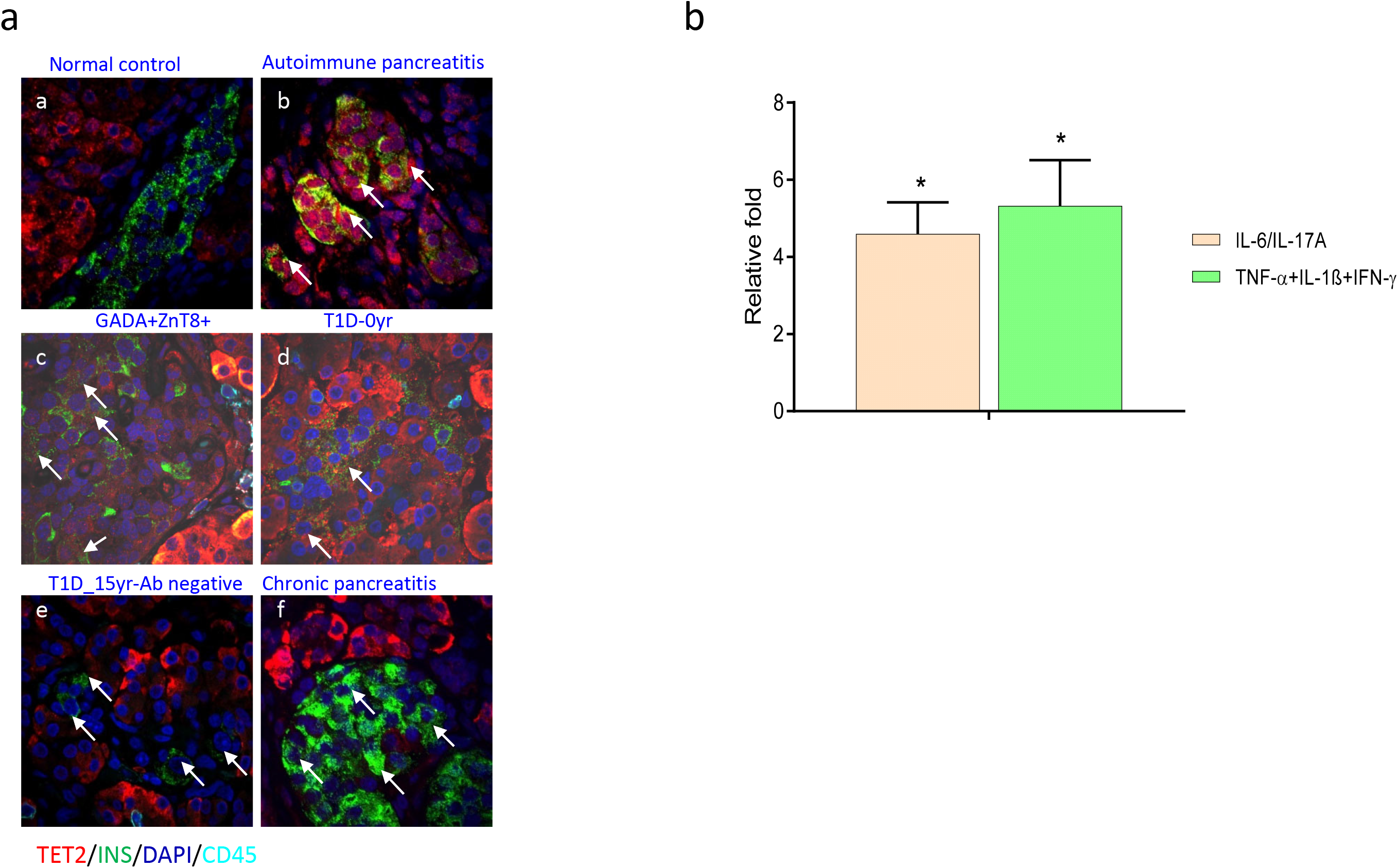
Induction of TET2 in human β cells with inflammation. (a) Expression of TET2 protein in human β cells in vivo in pancreas from a healthy control subject (a) and patient with autoimmune pancreatitis (b), a non-diabetic individual with autoantibodies to GADA and ZnT8 who is a relative of a patient with Type 1 diabetes (c), and a patient with recent onset T1D (d), patient with Type 1 diabetes for 15 years without detectable autoantibodies and (f) a patient with chronic pancreatitis. The sections were stained with anti-TET2 (red) and anti-INSULIN (green) as well as anti-CD45 (Cyan) antibody. DAPI (blue) stains cell nucleus. The merged staining is shown. Arrows indicate TET2+β cells in b, c and d. and β cells in e and f. Data represent normal individuals (*n*=8), donors with autoimmune pancreatitis (*n*=5), non-diabetic donors who were autoantibody+ (*n*=7), and C-peptide+ patients with T1D of relatively short duration (*n*=7). (b) Human islets were cultured with IL-6 (20 ng/ml)+IL-17A (50 ng/ml) or TNF+IL-1β+IFNγ (10 ng/ml each) for 24 hrs before FACS enrichment for Zn+TMRE+ <u>β</u> cells and transcription analysis of *TET2* gene by qRT-PCR. The fold of *TET2* mRNA induced by cytokines relative to media was shown. Data (mean ± SEM) are from 3 experiments, each with 2,000 islet equivalency (IEQ) from one non-diabetic donor (Fold vs media, one sample Student’s t-test vs media (*p=0.02).

Our findings were consistent with our studies with murine islets and suggested that inflammatory cytokines could induce *TET2* expression. We therefore analyzed *TET2* expression in human β cells from islets that were cultured with IL6/IL-17A or TNFα+IL-1β+IFNγ (**Fig. 2**b) (p=0.02 each) and found increased *TET2* gene expression.

These observations suggested that *TET2* expression was induced by inflammatory cytokines and suggested a permissive but not sufficient role for β cell killing since we found increased TET2 in β cells from patients with autoimmune pancreatitis that do not develop diabetes. However, we noted that not all of the β cells that were identified in our studied patients with T1D or in the autoantibody+ relatives were TET2+. Moreover, in samples from patients with long-standing T1D, we identified β cells by insulin staining, consistent with reports of residual β cells even in long-standing patients, but we did not identify TET2 expression in those β cells (**Fig. 2**a (e)).

### Resistance of Tet2-KO β cells to Immune Killing

To understand the role of Tet2 in the inflammatory responses of β cells, we bred Tet2-KO NOD mice. However, Tet proteins play an important role in cell development ^24^ and Dhawan et al. reported that deletion of Tet2 in the pancreatic lineage results in improved glucose tolerance and β cell function (https://doi.org/10.2337/db18-50-OR). Therefore, we first characterized β cell function, islet cells, and gene expression from Tet2-KO B6 mice. The glucose tolerance to IPGTTs and morphology of the islets were indistinguishable between the KO and wild type B6 mice (Supplementary **Fig. 1**a, b). There were small but significant differences in the proportion of β and α cells in the KO mice as well as in the MFI of Insulin and Pdx1 but the expression of *Ins1, Ins2, Nkx6.1, ChgA*, and *Pdx1* were similar to WT β cells (Supplementary **Fig 1**c-e). Overall these findings suggest that there may be subtle differences in the composition of islets from KO and WT mice but there were no detectable functional abnormalities of Tet2-KO β cells.

We backcrossed the KO allele to NOD for more than 14 generations. The Tet2-KO NOD mice did not develop spontaneous diabetes whereas the median time to diabetes was 17 and 23 weeks in their WT and HET littermates (Supplementary **Fig. 2**).

Because Tet2 is involved in the development and function of immune cells, we transplanted bone marrow from WT NOD mice into WT or Tet2-KO recipients to eliminate any impact of Tet2 loss in immune cells. The Tet2-KO mice had a markedly reduced incidence of diabetes (median survival undefined) compared to WT recipients of NOD WT BM (median survival =11 wks) (p<0.0001) (**Fig. 3**a). This was not due to the lack of immune infiltration in KO recipients. As shown in **Fig. 3**b, at the same level of immune cell infiltrates, there was a decline in the proportion of surviving β cells in the WT recipients (r=−0.82, p<0.0001) that was not seen in the Tet2-KO recipients (p=0.1447). It was possible that autoreactive cell development was impaired in the Tet2-KO recipients. However, we found a similar frequency of IGRP-reactive CD8+ T cells in the WT and Tet2-KO bone marrow recipients (**Fig. 3**c). In addition, we transplanted splenocytes from KO bone marrow transfer (BMT) recipients that did not have diabetes, or splenocytes from diabetic WT BMT recipients, into NOD/*scid* mice. Donor cells were from 11-14 week post-BMT mice. These two splenocyte-recipient groups of mice developed similar rates of diabetes (**Fig. 3**d), arguing that autoimmune T cells are present in non-diabetic Tet2-KO BMT recipients. To further understand the nature of the resistance of β cells to killing, we transplanted islets from 5-6-wk-old KO vs WT mice under the kidney capsules of 6-wk-old NOD WT recipients. 2 weeks after the transplant, all recipients were given 1 × 10^7^ diabetogenic splenocytes to synchronize the development of diabetes. Recipients were followed for the development of hyperglycemia which required killing of both the endogenous and the transplanted islets. The time to diabetes onset in the recipients of KO islets was delayed by 4.5 weeks (median survival=19.5 weeks) compared to recipients of NOD WT islets (median survival =15 weeks) (p<0.0001). All of the 11 recipients of WT islets developed diabetes whereas, at 25 weeks, 3 of the 12 recipients of KO islets were diabetes free (**Fig. 3**e). In these mice, there was rapid hyperglycemia following nephrectomy indicating that the transplanted islets were the source of insulin in the mice and that the endogenous WT β cells in the recipients had been destroyed.

**Fig. 3:**
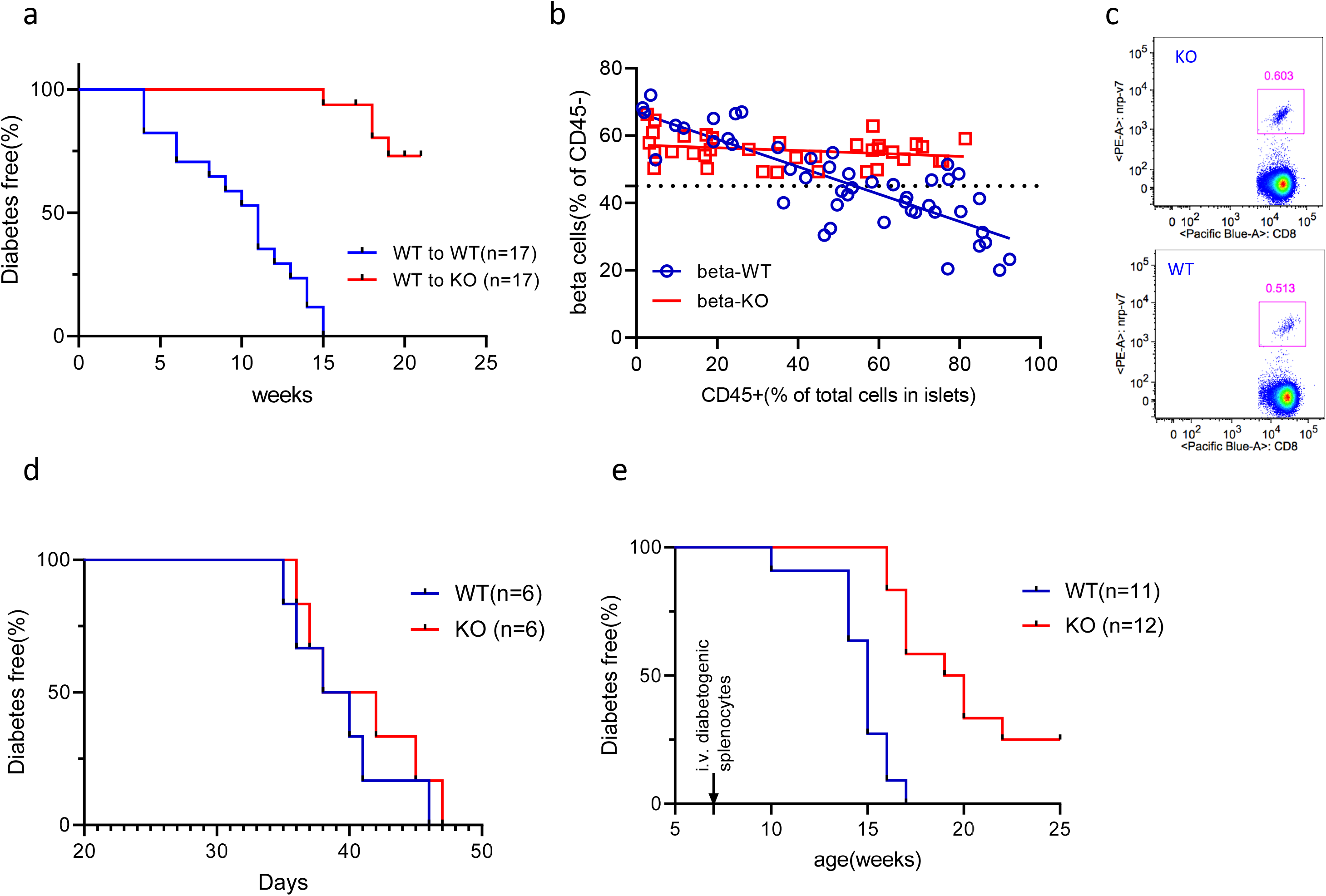
β cells are protected from immune killing in Tet2-KO NOD mice. (a) Diabetes incidence in Tet2-WT or KO NOD recipients of bone marrow from WT NOD mice. WT or Tet2-KO mice were lethally irradiated (total dose of 1100 rads) at 6-8 wks and were given 10^7^ bone marrow cells from 6-8-wk-old NOD mice. Recipients were followed for diabetes incidence up to 25 weeks (Log-rank curve comparison, p<0.0001, *n*=17 mice each group). (b) Relationship between cellular infiltrates and loss of β cells in WT (blue) and Tet2-KO recipients (red) of WT bone marrow. Shown here is the frequency of the intra-islets CD45+ cells and β cells (insulin+ by intracellular staining) analysed by FACS. Percentage of β cells was of CD45-islets cells. For WT recipients, r=−0.82, p<0.0001; for Tet2-KO recipients, p=ns. Each circle or square represents one recipient mouse (WT, n=47; KO, n=36). (c) Frequency of IGRP-reactive CD8+ T cells in the spleen from bone marrow recipients analysed by FACS. 14 weeks after transplant, splenocytes from KO or WT recipients of WT bone marrow were stained for IGRP-reactive CD8+ T cells with NRPV7 tetramer. A single pair of recipients of the same bone marrow are shown. Data represent at least 3 independent experiments with one pair of KO vs WT recipients per experiment. (d) Transfer of diabetes into NOD/*scid* mice that were given 10^7^ splenocytes from either KO BMT recipients that were not diabetic or WT BMT recipients that were hyperglycemic 12 weeks post-BMT. Data are from 2 experiments, each with pooled splenocytes from 2 KO and 2 WT BMT recipients (Log-rank curve comparison, p=ns). (e) Diabetes incidence in islet transplant recipients from either WT or Tet2-KO NOD islets donors. 250-300 pooled islets from 5-6-week-old WT or KO NOD mice were transplanted under the kidney capsule of 6-week-old WT NOD recipients. Two weeks post the transplant, all recipients received 10^7^ diabetogenic splenocytes. Blood glucose was measured twice weekly and diabetes incidence was recorded at the age of recipients. The median survival time in WT is 15 weeks vs 19.5 weeks in KO recipients. (Log-rank curve comparison, p<0.0001. WT, n=11; KO, n=12).

The resistance of KO β cells to killing was not absolute, since the Tet2-KO β cells could be killed by chemical means (i.e. streptozocin) in vivo (Supplementary **Fig. 3**) suggesting that the resistance of the KO β cells to killing was involved with immune mechanisms.

To directly assess the effects of inflammatory mediators on Tet2-KO β cells, we cultured singular islet cells with inflammatory cytokines that cause β cell death and found increased survival of Tet2-KO vs WT β cells (**Fig. 4**a). The differences in the WT and KO islets cells was seen in β but also, to a lesser degree in α cells (**Fig. 4**a). Second, we cultured WT NOD/*scid* and Tet2-KO β cells with purified islets infiltrates from pre-diabetic WT NOD mice and found a significant reduction in the frequency of live β cells when the WT vs Tet2-KO β cells were cultured with the diabetogenic immune cells (**Fig. 4**b).

**Fig. 4:**
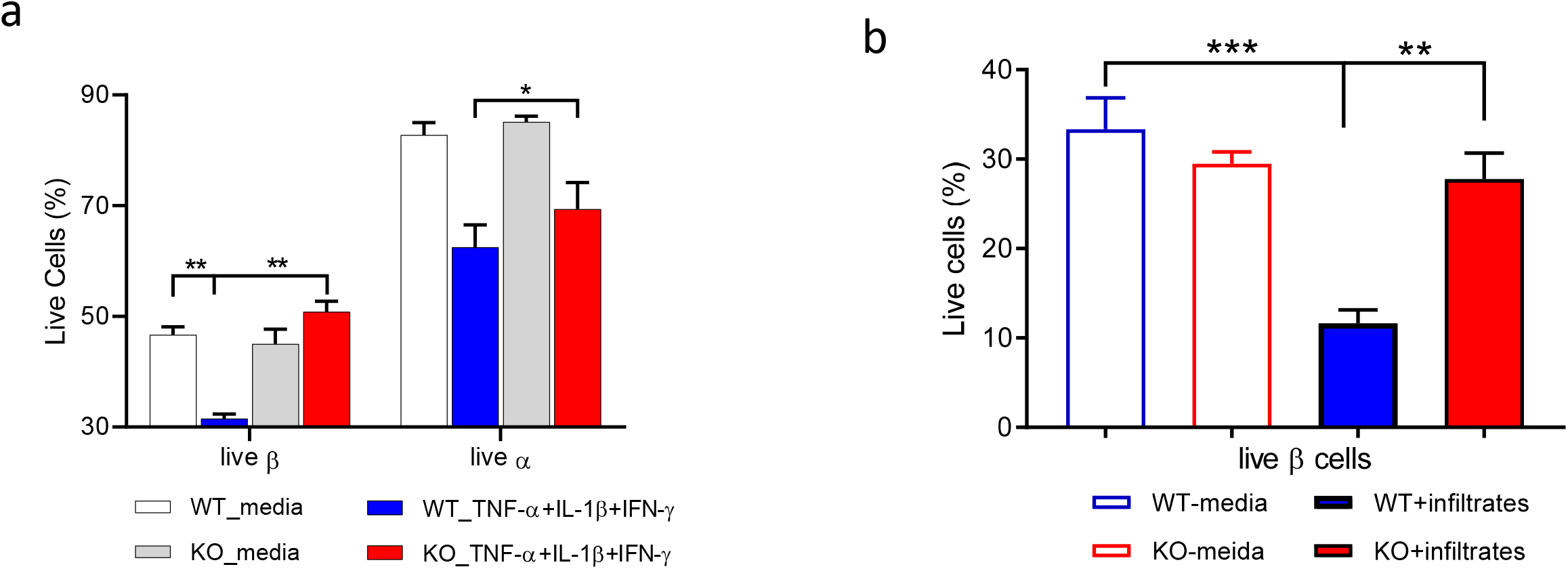
β cells lacking Tet2 are protected from immune-mediated killing in vitro. (a) Islet β cell survival during culture with cytokines. Single islet cells from WT or Tet2-KO B6 mice were cultured with the indicated cytokines for 12 hrs and the live cell percentage was determined in β as well as α cells (identified by insulin and glucagon staining) by FACS. Results are mean ± SEM of 3 experiments, *n*=6 mice per group per experiment (\**p*<0.05, \*\**p*<0.01, ANOVA). (b) β cell survival during culture with diabetogenic islets infiltrates. Sorted β cells from either WT NOD/*scid* (blue) or Tet2-KO NOD mice (red) were cultured overnight with islets infiltrates sorted from the same pre-diabetic WT NOD donors at a ratio of 1:5. Results are mean ± SEM of 3 experiments, *n*=5-6 mice per experiment (**p<0.0001, ***p<0.001, ANOVA).

These initial observations suggested that a subpopulation of β cells that survive immune attack have reduced Tet2 despite the general increase in expression that we had observed during disease progression and in human β cells that were exposed to inflammatory cytokines. In our previous studies in NOD mice, we identified a subpopulation of β cells, characterized by reduced insulin granularity, increased expression of “stem-like” genes and reduced β cell markers such as *Pdx1*, *Nkx6.1* and *MafA* that was resistant to immune mediated killing. Therefore, we analyzed Tet2 expression among these subpopulations of β cells from NOD mice. We found that Tet2 levels were higher in the normal (i.e. “top”) β cells that we showed succumbed to immune killing during progression of the autoimmune disease when compared to the hypogranular (i.e. “bottom”) cells that survived cytokine and immune cell mediated killing (Supplementary **Fig. 4**).

### Immune Responses are Not Activated by Tet2 Deficient Islet Cells

We analyzed the islet infiltrates from the WT and KO recipients of WT bone marrow, 8 weeks after transplant to understand how the Tet2-KO islet cells changed the immune responses,. The frequency of infiltrating cells was similar in the KO and WT islets (Supplementary **Fig. 5**). We analyzed gene expression by NanoString (pan-immunology panel) and found differential expression of 189 of 770 genes. There was decreased expression of genes associated kinase signaling needed for T cell activation such as Stat3, Jak2, Smad3, Nfkb, Mapk, reduced expression of molecules associated with cell-cell interaction as well as T cell function such as Cd137, Gzmm and Gzmk in addition to a panel of reduced cytokines (**Fig. 5**a). In addition, the infiltrating cells from KO recipients show increased expression of pro-cell death genes such as Bax, Casp3, Casp8 and Tnfsf10 (Trail) and reduced pro-survival gene Bcl2. Pathway analysis (with Ingenuity Pathway Analysis (IPA)) identified differences in inflammatory responses implicating 91 genes. Other affected pathways included cell-to-cell signaling and interaction, cellular function and mainenance, growth and proliferation and movement which were significantly decreased in immune cells from KO recipients. There were phenotypic difference between the infiltrating T cells in the KO and WT mice. Th1 and Th2 activation pathway were reduced in KO recipients as well as pathways of cell trafficking, (**Table 1**). To confirm these findings, we analyzed immune cells from the draining (pancreatic) lymph nodes from KO and WT BMT recipients of WT bone marrow by FACS. The T cells from the KO recipients showed reduced frequencies of CCR7+ and CXCR3+ CD4 T and CD8 T cells and fewer central memory CD4+ T cells (**Fig. 5**b).

**Fig. 5:**
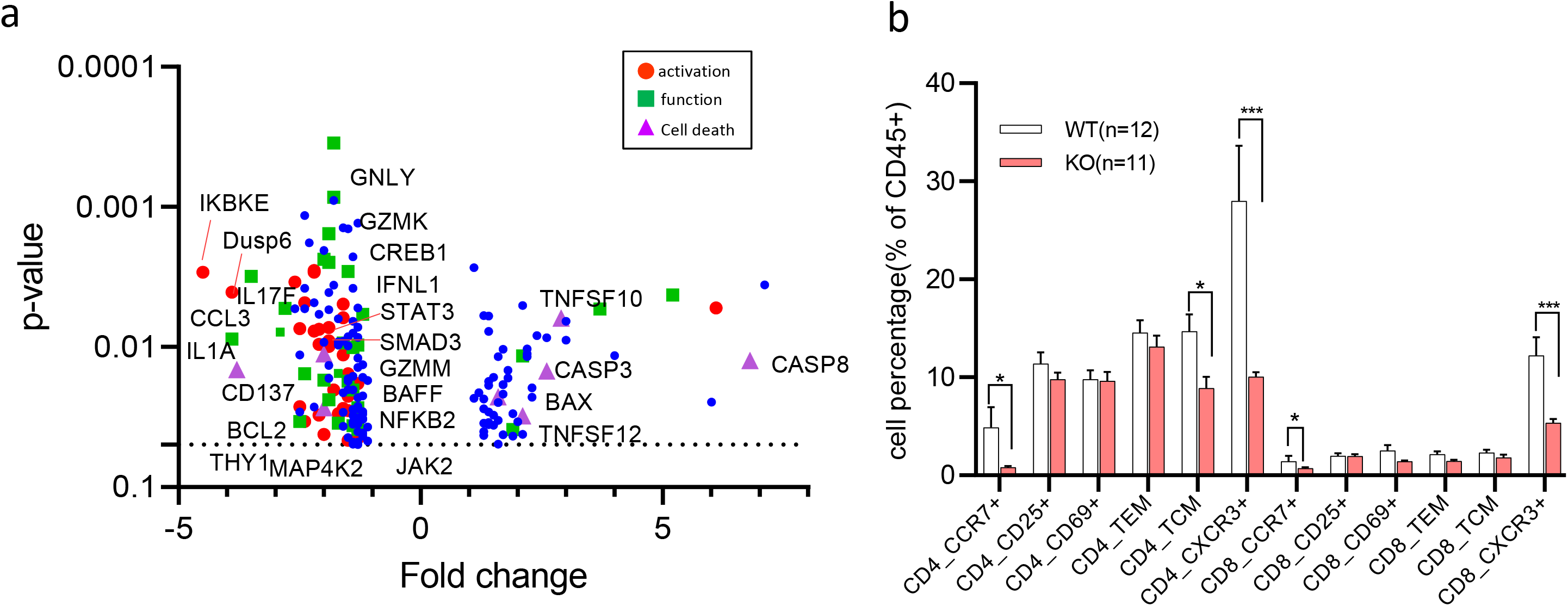
T cells from Tet2-KO islets are less activated and show reduced pathogenic phenotypes. (a) Volcano plot showing transcription differences in islets infiltrates between KO and WT recipients of WT bone marrow. 8 weeks post-BMT, CD45+ islets infiltrates were sorted from KO vs WT recipients and used subsequently for transcription analysis with NanoString PanImmunology panel. 189 of 770 genes that were found to be different between infiltrates from WT and KO recipients are shown (after FDR correction, p<0.05). Genes that are associated with T cell activation, cell function as well as cell death are highlighted with different colors among the 189 genes. Data are from 6 KO vs 6 WT samples that were pooled from 9 KO and 8 WT mice respectively. (b) FACS analysis of T cells from pLNs of KO vs WT recipients of bone marrow from WT NOD mice. The % of CD45+ cells is shown. TCM=CD44+CD62L+. Data are from 4-5 experiments with pLNs from an individual mouse as one sample (mean ± SEM) (WT: *n*=12; KO: *n*=11).

**Table 1:** Diseases and Bio Functions identified by Ingenuity Pathway Analysis of Nanostring data. Diseases and Bio Functions identified by Ingenuity Pathway Analysis of PanCancer Immune Profiling Data. CD45+ infiltrates were sorted from KO vs WT recipients 8-weeks post-BMT from WT NOD bone marrow donors (n=13-16 KO or WT mice over 6 separate sortings). 9,000-45,000 cells were lysed for each reaction. Results were summarized into different disease and Bio function (marked in bold). The name of specific pathway was listed as well as the P value and number of molecules involved in each pathway.

|  |  |  |
| --- | --- | --- |
| **Diseases and Disorders** | p-value range | #Molecules |
| Inflammatory Response | 1.22E-04 – 9.69E-49 | 91 |
| Immunological Disease | 8.60E-05 – 2.59E-26 | 44 |
| **Molecular and Cellular Functions** | p-value range | #Molecules |
| Cellular-To-Cell Signaling and Interaction | 1.22E-04 – 9.69E-49 | 86 |
| Cellular Function and Maintenance | 1.22E-04 – 1.43E-45 | 85 |
| Cellular development | 1.22E-04 – 1.58E-42 | 82 |
| Cellular Growth and Proliferation | 1.22E-04 – 1.58E-42 | 84 |
| Cellular Movement | 1.22E-04 – 8.71E-41 | 62 |
| **Physiological System Development and Function** | p-value range | #Molecules |
| Immune Cell Trafficking | 1.22E-04 - 9.69E-49 | 90 |
| Lymphoid Tissue Structure and Development | 1.22E-04 - 1.58E-42 | 92 |
| **Top Canonical Pathways** | p-value | Overlap |
| Th1 and Th2 Activation Pathway | 9.14E-21 | 15.2 % 23/151 |
| Th1 Pathway | 8.92E-19 | 18.1 % 19/105 |

### β Cells Lacking Tet2 Show Reduced Inflammatory Responses

The differences in WT and KO β cells might affect the infiltrating immune cells. Therefore we sorted β cells (Zn+TMRE+) from the WT or KO recipients of WT bone marrow, 8 weeks after WT transplant and analyzed the cells by bulk RNA-seq. 333 genes were found to be differentially expressed between KO and WT β cells (p < 0.05 after FDR correction (Supplementary **Fig. 6**)), and 199 of these genes showed ≥1.5 fold change in expression (**Fig. 6**a, b). We identified a number of differences in pathways associated with β cell death and immune protection. These included reduced PTEN signaling (p=0.00129) ^25,26^, inflammatory mediators (*Irf8, Tifa*), attenuated immune signaling (*Pik3cd, Abl2*) and cytokine inducible genes (*Gbp2, Gbp6, Ifi47, Tnfaip2*). Genes involved in Class I and II MHC expression were reduced but the levels of Class I MHC were not reduced on β cells from the Tet2-KO mice (Supplementary **Fig. 7**). Meanwhile, β cells from KO recipients expressed genes associated with immune suppression (GABA receptor A (Gabrg3) ^27^, Gad2, CD14 ^28^and H2-Q7 (Qa-2) ^12 13^ (**Fig. 6**a, b). Compare to WT β cells, there are also changes in KO β cells that are related to improved β cell fitness (*Slc39a4, Tgfbr2, Fgfr2, Fgfr3 and Aldh1a3*) ^29 30–32^ as well as improved β cell survival (*Gas6, Ptpn13, Tnfrsf19*) ^33 34 35–37^ (**Fig. 6**a, b).

**Fig. 6:**
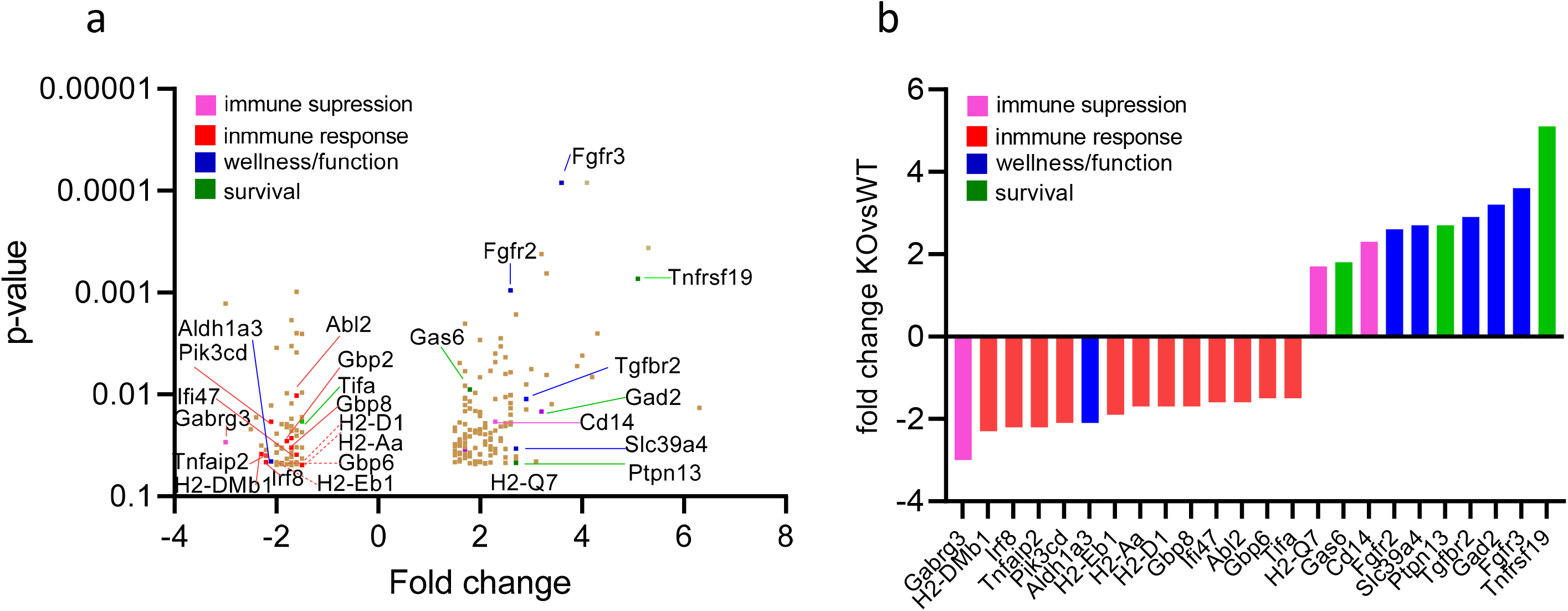
β cells lacking Tet2 have reduced inflammatory responses. (a) Volcano plot showing gene expression differences in β cells from KO vs WT recipients of WT bone marrow. 8 weeks after the bone marrow transfer, β cells that are TMRE+Zinc+ were sorted from KO vs WT recipients and subjected to cDNA library preparation and bulk RNA-seq analysis. There are 333 differentially expressed genes in total with p<0.05 after FDR correction, and 199 with fold change>1.5 that are shown in A. Genes of particular interest are color coded in a and b according to their function. Data are from 3 sortings and 3-4 mice per group per sorting.

To understand the basis for the epigenetic control of the inflammatory responses in β cells, we performed ATAC-seq on the same β cell samples analyzed by RNA-seq (n=2 KO, 3 WT). We identified 60,449 open chromatin sites among the five samples profiled. Accessibility at the overwhelming majority of these sites (~97%) did not differ substantially between WT and Tet2 KO cells, suggesting that Tet2 deficiency does not lead to widespread chromatin remodeling of ß cells. 715 sites (~1%) had greater accessibility in KO’s (FC > 1.5) and 1387 (~2%) had lower accessibility (FC < −1.5) in Tet2 KO vs. WT cells (**Fig. 7**a). Using HOMER transcription factor (TF) motif enrichment analysis, 15 TF motifs were identified as enriched in peaks with higher accessibility (opening peaks) in KO β cells (q-value < 0.05) (Supplementary **Table 2**). Most of these corresponded to TFs controlling islet β cell identity and function such as Foxa1, Foxa2 and Rfx and were also enriched in closing peaks in KO β cells (Supplementary **Table 1**), suggesting that Tet2 deficiency results in opening and closing of a subset of islet TF binding sites. 97 TF motifs were enriched in peaks with lower accessibility (closing peaks) in KO β cells (q-value < 0.05) (Supplementary **Table 1**). Interestingly, motifs for 15 TFs mediating inflammatory responses such as Stats 1, 3, 4, and 5, and Irfs 1, 2, 4, and 8 (**Fig. 7**b) were specifically enriched in closing peaks, suggesting that inflammatory stress-responsive *cis*-regulatory elements are epigenetically decommissioned in Tet2 deficient ß cells.

**Fig. 7:**
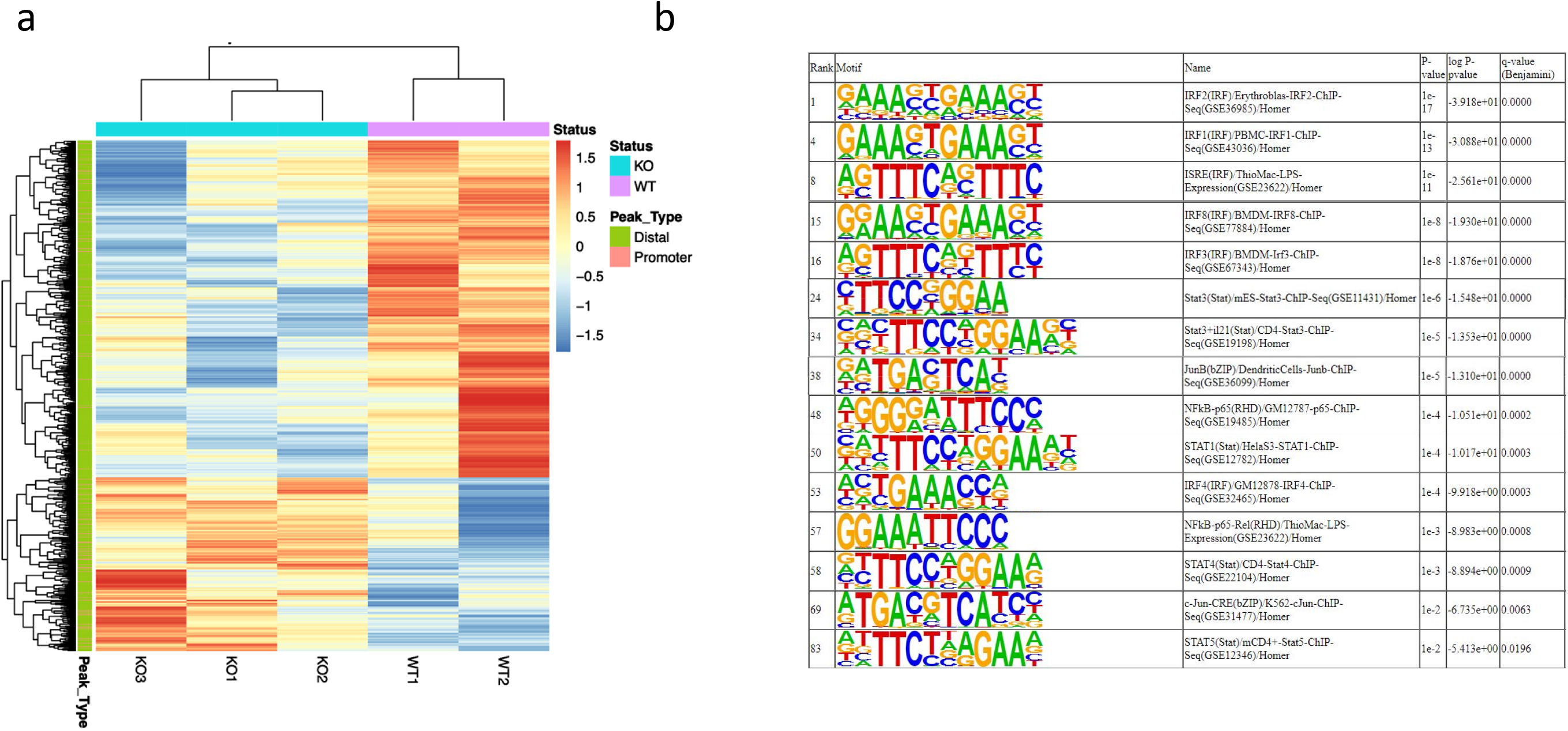
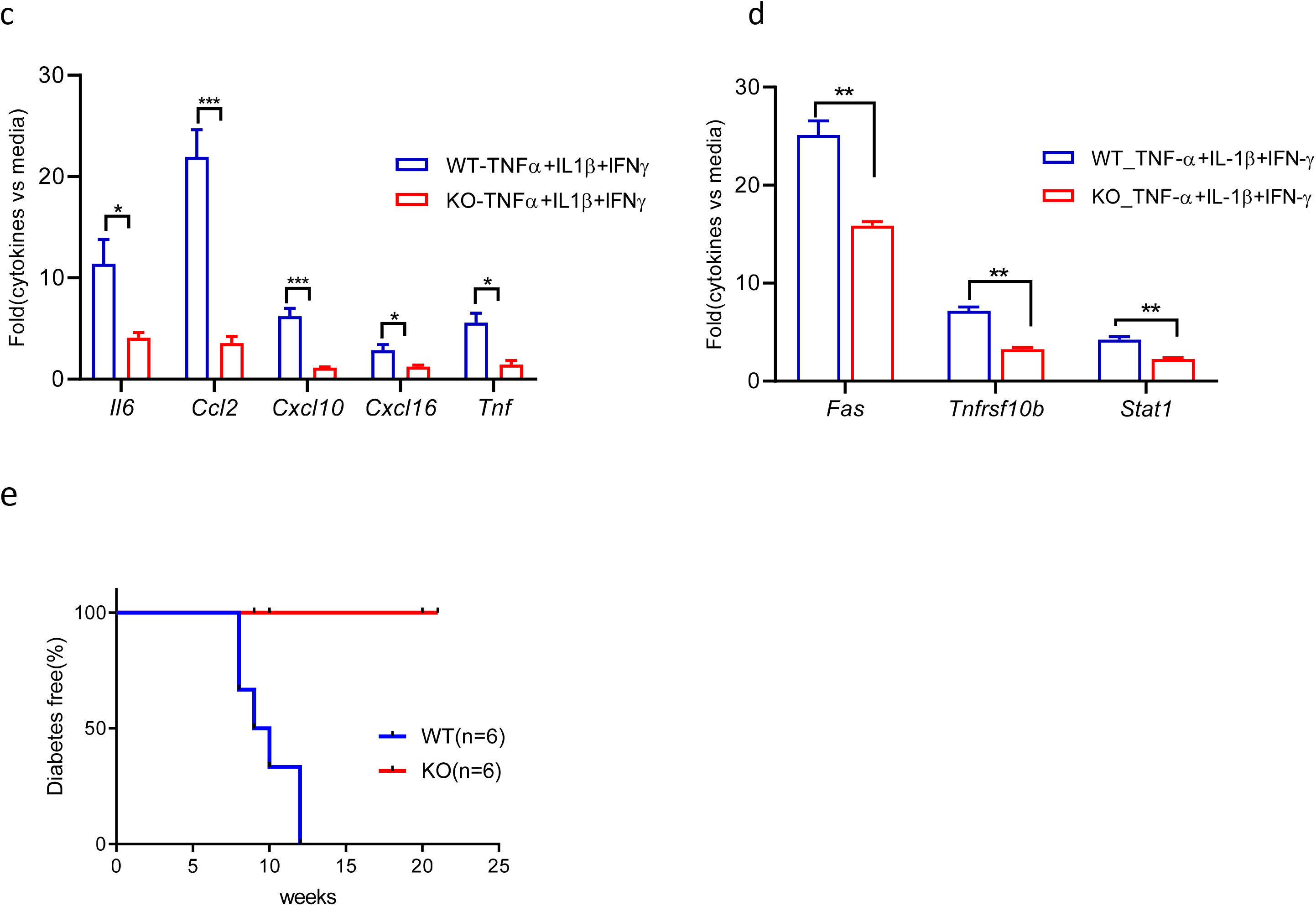
Tet2-KO ß cells show reduced chromatin accessibility at putative inflammatory mediator binding sites, reduced inflammation response, and are resistant to killing by transferred diabetogenic cells. (a) Comparison of <u>A</u>TAC-seq profiles identify chromatin accessibility changes in β cells from KO vs WT recipients of WT bone marrow. Of 60,449 accessible sites identified across the five samples, 715 (~1%) had increased accessibility (FC > 1.5) and 1387 (~2%) had reduced accessibility in (FC < −1.5) in KO’s, respectively. (b) HOMER motif analysis identified 97 TF motifs enriched in peaks with decreased accessibility (closing peaks) in KO β cell samples (q-value < 0.05), 15 of which are inflammatory mediators. The rank of the 97 TF motifs was sorted by q-value. (c, d) transcription analysis of cytokines and chemokines (c) as well as cell death receptors (d) in KO vs WT islets following culture with the indicated cytokines for 24 hrs. Data are mean ± SEM of 4 independent experiment, 4 mice per group per experiment (\**p*<0.05, **p<0.0001, ***p<0.001, ANOVA). (e) Diabetes incidence in Tet2-WT or KO NOD mice that received by adoptive transfer 1.5 × 10^7^ diabetogenic splenocytes from diabetic WT NOD mice after half lethal dose irradiation (650 rads). Mice were followed up to 21 weeks post adoptive transfer (Log-rank curve comparison, p=0.0041, n=6 each group).

To confirm these findings and the responses of β cells to inflammatory mediators, we first screened inducible cytokine and chemokine species in islets following IFNγ with or without TNFα+IL-1β. *Il6, Cxcl10, Cxcl16, Ccl2,* as well as *Tnf* can be induced by this cytokine cocktail (TNFα+IL-1β+IFNγ) but not C*cl12, Ccl19* or *Il1b* (Supplementary **Fig. 8**a).

Previously Xu et al described impaired STAT1 signaling in tumor cells cultured with IFNγ ^19^ but we did not identify any difference in IFNγ response measured by the production of chemokines or expression of *Pdl1*or *Ido in* KO vs WT islets from either NOD (Supplementary **Fig. 8**b) or B6 mice (Supplementary **Fig. 8**c).We cultured WT and KO islet cells with TNFα+IL-1β+IFNγ and found in Tet2-KO islets reduced induction of *Il6, Ccl2, Cxcl10, Cxcl16* and *Tnf,* which have been linked to β cell destructive processes (**Fig. 7**c). In addition, we found that the expression of genes associated with β cell death such as *Fas, Tnfsf10b* (TRAIL-R2) and *Stat1* ^38,39^ were also reduced in cultured KO β cells (**Fig. 7**d)

Finally, to determine whether these differences in inflammatory responses are associated with β cell survival, we transferred splenocytes from diabetic NOD mice into irradiated WT and KO recipients and followed them for the development of diabetes. The median time to hyperglycemia was 9.5 weeks in the WT recipients whereas none of the 6 KO recipients developed diabetes for more than 20 weeks after adoptive transfer (p=0.0041) (**Fig. 7**e)

## Discussion

We have found that TET2 can control immune mediated destruction of human and murine β cells through the interactions with immune cells and cell intrinsic mechanisms. *Tet2* expression is increased during progression of diabetes in islet cells and in β cells. However, a subpopulation of β cells that we previously showed was protected from autoimmune killing has lower levels of *Tet2* expression. In human pancreases, β cells also express *TET2* under inflammatory conditions, but islets from normal individuals and from patients with long-standing T1D have β cells without detectable *TET2* expression. Our findings indicate that *TET2* is associated with, but not sufficient for, killing of β cells, since we also found increased *TET2* expression in β cells from patients with autoimmune pancreatitis who do not develop diabetes. Instead, our studies in Tet2-KO mice indicate that in the absence of Tet2, β cells are protected from immune-mediated killing. We used three complementary model systems: bone marrow transplantation, islet transplantation, and adoptive transfer of diabetogenic splenocytes to identify the interactive relationships between Tet2 expression in β cells and immune cells and show that Tet2-KO islets are resistant to killing even in the presence or transfer of diabetogenic T cells. The Tet2-KO β cells show transcriptional differences in response to inflammatory cytokines that involve increased expression of genes associated with β cell wellness, function, survival as well as genes for immune suppression, and reduced expression of genes associated with immune responses and activation. Importantly, there was also reduced chromatin accessibility at putative regulatory elements containing binding motifs of inflammatory mediators in Tet2-KO β cells. These molecular changes were confirmed by functional studies that showed reduced secretion of mediators when the Tet2-KO islets were cultured with inflammatory cytokines. As a result, Tet2-KO islets fail to activate immune responses and autoreactive T cells, resulting in reduced autoimmune elimination of Tet2-KO ß cells compared to WT. Our studies have identified a mechanism that controls interactions between β cells and immune cells that leads to immune-mediated killing. Future studies to inhibit this pathway may lead to improved survival of β cells in the setting of autoimmunity.

In human pancreatic sections, we found differences in TET2 expression within islet cells and between pancreases from individuals with different stages of β cell destruction. We found expression in both the nuclear and cytoplasm, consistent with reports in neurons ^40^. In patients with long standing T1D, the absence of TET2 expression suggests an association with survival. Our studies of the control of β cell responses by Tet2 provide insight into the mechanisms that account for these histologic findings. A fully pathologic autoimmune repertoire developed in the Tet2-KO recipients of WT bone marrow. However, these cells did not receive the same activation signals and did not cause β cell killing as shown by our analysis of their transcriptome when compared to islet infiltrates from the WT BMT recipients. The absence of TET2 expression on β cells, therefore, could potentially explain the ability of some β cells to survive even when autoreactive T cells remain in the host. Nonetheless, when T cells from the Tet2-KO bone marrow recipients were transferred to WT NOD/*scid* recipients, diabetes occurred at a similar rate. Consistent with these findings, islets from the Tet2-KO mice showed significantly delayed rejection compare to islets from WT donors when they were transplanted into WT NOD recipients. Even in the same mouse, the WT but not KO islets were preferentially destroyed.

These observations indicate that the islet cells themselves deliver signals to the autoreactive immune cells that leads to their activation and β cell killing. These signals most likely are the result of differences in the Tet2-KO β cells since the antigen presenting cells in the transplant recipients were from the WT bone marrow donor in the case of BMT or WT NOD mice in the case of islets transplant. Release of inflammatory cytokines, which is well documented to occur from β cells may account for the difference. For example, activation of pathologic T cells has been shown to be induced by CXCL10 that binds to CXCR3 on the surfaces of Th1 cells ^41^.

ATAC-seq revealed reduced chromatin accessibility in Tet2-deficient ß cells at putative binding sites for Stat and Irf TFs that have been associated with cytokine signaling. Previous studies of Tet2-deficient tumor cells identified impaired STAT1 signaling that prevented the expression of chemokines and PD-L1 expression and enabled tumors to evade anti-PD-L1 immune therapy ^19^. These investigators described a disrupted IFNγ-JAK-STAT signaling cascade in Tet2-deficient tumor cells but we did not see the same with β cells.

In addition to the participation of the β cells in immune cell interactions, we also identified pathways to account for enhanced β cell survival in Tet2-KO β cells. Our culture studies identified improved survival when β cells were cultured with inflammatory cytokines as well as infiltrating immune cells, both of which may mediate β cell killing. In the Tet2-KO β cells, expression of GABA receptor A (*Gabrg3*), which is associated with immune suppression, was reduced and there was increased expression of *Tnfrsf19, Fgfr2 and Fgfr3*. Increased *Tnfrsf19* expression was shown to attenuate TNFβR signaling and improve β cell survival ^29 30^. Fgfr2 and Fgfr3 are growth factor receptors, and attenuation of FGF signaling by expressing dominant-negative forms of the FGFRs receptors in mouse β cells leads to diabetes ^31^. Gas6, which is linked to β cell proliferation, was also increased in expression on the KO β cells ^42^. These cell-intrinsic mechanisms may account for the resistance of the cells to killing by inflammatory cytokines in the absence of extrinsic cells.

There are a number of limitations and unresolved aspects of our studies. Tet2 causes demethylation of methylcytosines, and therefore the relationship between the loss of this enzymatic activity and decreased transcription of genes is somewhat paradoxical. However, our previous studies showing increased activity of Dnmts suggest a mechanism that, if unopposed, could lead to these outcomes ^43^. In addition, the relationship, particularly in human tissues, between the autoreactive repertoire, TET2 expression and β cell killing will require further studies with human β and effector T cells. Finally, our murine studies involve complete elimination of Tet2 from islet cells.

In summary, our studies have identified a novel interaction between β cells and immune cells that regulates immune cell activation, recruitment and killing. Expression of Tet2 is required for β cells to produce chemokines and cytokines that are needed for the activation and pathologic function of diabetogenic T cells. In its absence, β cells resist cell- or cytokine-mediated killing even when autoreactive cells are present through cell intrinsic mechanisms as well as avoiding activation of immune cells. These studies suggest a future strategy to protect insulin producing cells from immune-mediated killing in the setting of autoimmunity or even for engineering more resilient and/or resistant β cell replacements to avoid recurrent immune elimination.

## Methods

### Mice

TET2-KO male breeder under B6 background was purchased from The Jackson lab and backcrossed with NOD females for over 14 generations. Female NOD, B6, and NOD/*scid* mice were also obtained from The Jackson Laboratory and maintained under pathogen-free conditions. All protocols were approved by the Yale Institutional Animal Care and Use Committee.

### Human Islets and Pancreas Tissues

Human islet samples were obtained from adult, non-diabetic organ donors from the Integrated Islet Distribution Program. Pancreas tissues from donors with autoimmune pancreatitis (n=5) or chronic pancreatitis (n=3) were obtained from the Department of Pathology at Yale and pancreas slides from nPOD include normal individuals (n=8), non-diabetic donors who were autoantibody+ (n=7), and C-peptide+ patients with T1D of relatively short duration (n=7). The use of human tissues was approved by the Yale Institutional Review Board.

### Immunofluorescent labeling of human pancreas

To visualize TET2 protein expression in pancreatic islet cells, formalin-fixed, paraffin-embedded sections (5μm thickness) of human pancreas were incubated with primary antibodies against TET2 (Abcam, cat# 230358), INSULIN (DAKO, cat# A0564) and CD45 (DAKO, clone# 2B11 + PD7/26) overnight at 4 °C, washed and processed with the appropriate secondary fluorescent-conjugated antibodies. Subsequently, sections were dyed with 0.7% Sudan Black and CuSO4 to quench auto-fluorescence and counterstained with DAPI, as previously described ^44^. Images were acquired on an UltraVIEW VoX (PerkinElmer) spinning disc confocal Nikon Ti-E Eclipse microscope using the Volocity 6.3 software (Improvision).

### Mouse Islet Isolation and β Cell Staining and Purification

Mouse islets were handpicked with a stereomicroscope after collagenase digestion of pancreases (^15^. Single-cell suspensions were prepared as previously described (^13^). For β cells enrichment, single islet cells were stained with TMRE and FluoZin-3 and sorted with a FACSAria II (BD). In some experiments intracellular staining with anti-insulin (R&D Systems, Clone # 182410), anti-Tet2 (Cell Signaling Technologies, clone# D6C7K) and anti-glucagon (Abcam, clone# K79bB10) was done to identify β cells, α cells that was analyzed or sorted with LSRFortessa (BD).

### Islet Cultures

Human or mouse islets were rested upon arrival or isolation in complete RPMI media for 3 hours before aliquoting (100 per well) into a non-treated 24-well plate and culture for 18-24hrs with or without cytokines. The cells were then harvested for RNA extraction and transcriptional analysis of genes of interest. In cytokine or immune cell mediated killing experiments, mouse islets were dispersed with accutase and cultured as single cells overnight before harvest for live/dead analysis by flow or cell death receptors analysis by qRT-PCR.

### Adoptive Transfer of Diabetogenic Splenocytes

Tet2-KO or WT NOD mice (8 weeks of age) were irradiated with split (2 × 325 rads, 2–3 h apart, 650 rads total) dose and splenocytes (1 × 10^7) from a WT NOD mouse with recent onset DM were transferred i.p. within 3 h of irradiation. Diabetic incidence was followed and recorded as the time (weeks) post-adoptive transfer.

### Streptozocin (STZ) treatment

8 to 10-wk-old Tet2-WTand KO B6 mice were given a single dose of STZ (200 mg/kg, i.p) and followed daily for diabetes following the treatment.

### Bone Marrow Transfer (BMT)

Bone marrow transfer (BMT) from and to NOD mice were done as previously described ^45^. Donor NOD mice 6–8 weeks of age were euthanised by CO2. Bone marrow (BM) was flushed with ice cold HBSS and erythrocytes removed by lysis. For BMT, recipient KO or WT mice (4–8 weeks of age) were irradiated with split (2 × 550 rads, 2–3 h apart, 1100 rads total) dose and donor BM (10*10^6 cells) transferred i.v. within 3 h of irradiation Recipients were followed for hyperglycemia (glucose > 250 mg/dl × 2) and diabetic incidence was recorded at the time (weeks) post-BMT.

### Islet Transplant

250-300 hand-picked islets were pooled from 5-6 weeks KO or WT donor (2-3 donors per recipient). The islets were then put under the kidney capsule of WT NOD recipients (6-weeks of age). 2 weeks later, diabetogenic splenocytes were given via i.p.to islets transplant recipients (1x 10^7 cells per recipient). Blood glucose was measured twice weekly and diabetes incidence was followed and recorded at the age of islets recipients (weeks).

### Intraperitoneal Glucose Tolerance Test (IPGTT)

B6 Tet2-WT, HET and KO mice were fasted for 16 hr. Blood glucose levels were measured 0, 15, 30, 60, and 120 min after glucose injection (2 g/kg body weight). IPGTT result was presented as area under the curve using trapezoidal rule ^46^ and then divide by the total assay time (120 min).

### Low-Input RNA-Seq and Data Analysis

Viable beta cells were enriched by FACS sorting from KO or WT recipients 8 weeks post-BMT from WT bone marrow donors. RNA was extracted and reverse transcription and cDNA amplification were performed using 10 ng of RNA using the SMARTer Ultra Low RNA kit (Clontech Laboratories) like previously reported ^13^. Sequencing libraries were prepared using Nextera XT DNA Sample Preparation kit (Illumina). Libraries were pooled and sequenced on an Illumina HiSeq4000 using single-end 100-bp reads. This sequencing service was conducted at Yale Stem Cell Center Genomics Core facility which was supported by the Connecticut Regenerative Medicine Research Fund and the Li Ka Shing Foundation. Mapping of reads to mouse reference transcriptome mm9 and quantification of mRNA expression. RNA-Seq data were analyzed with Partek Flow software (v6.6).

### NanoString Pan-Immunology Panel Analysis of Islet Infiltrates

Analysis was done according to nanoString technologies user manual for nCounter XT CodeSet Gene Expression assays. Briefly, 9,000-45,000 CD45+ islets infiltrates were sorted from KO vs WT BMT recipients from WT NOD and cell pellets were lysed in 33% RLT lysis buffer from QIAGEN and up to 3.5μl lysate was used directly in a 15μl hybridization reaction which was kept at 65°C for 24 hrs. The hybridization product would then been cleaned up on automatic Prep Station, after which the end product would be ready for the digital analyzer nCounter MAX/FLEX system which rapidly immobilizes and counts samples which have hybridized to nCounter barcodes. The raw data were then downloaded from the digital analyzer and further analyzed through nSolver4.0 analysis software, also provided also by nanoString technologies. Differentially expressed genes (P value < 0.05) were used for functional analysis with Ingenuity Pathway Analysis and upstream regulator analysis (https://www.qiagenbioinformatics.com/products/ingenuity-pathway-analysis/).

### RNA extraction and cDNA synthesis

For fresh, cultured as well as FACS-sorted cells, RNeasy Plus Mini Kit (QIAGEN, cat#74134) was used for RNA extraction and High-Capacity cDNA Reverse Transcription Kit was used for cDNA synthesis (Applied Biosystems, cat#4368814).

### Probes, Primers, and Real-Time qPCR

Primer pairs were used in this study together with a QuantiFast SYBR green PCR kit (QIAGEN). The sequences of primer pairs used in this study are listed in Supplementary Table 3. The *Actb* housekeeping gene was used to normalize the input RNA in all real-time qPCR assays. Gene transcription was presented as ΔCt = (Ct of Actb -Ct of target gene) + 20 or in other cases as relative fold to control.

### ATAC-seq and Data Analysis

FACS sorted mouse islet β cells ATAC-seq libraries were prepared as previously described ^47^. After wash and lysis, cell pellet was resuspended in 50 ul of transposition mixture (Illumina Tagment DNA TDE1 Enzyme and Buffer) and incubated 30 min at 37°C. Transposed DNA was cleaned up over Qiagen MinElute kit and stored frozen at −20°C until indexing following the protocol in Ref. (^48^. Libraries were sequenced on an Illumina NovaSeq S1 with 2 × 101 bp cycles. Raw sequence reads were quality trimmed using Trimmomatic version 0.32 ^49^ with the following parameters “TRAILING:3 SLIDINGWINDOW:4:15 MINLEN:36.” Trimmed reads were aligned to mouse genome (mm10) using BWA version 0.7.12 ^50^, specifically using the bwa mem –M option. Duplicate reads were marked and removed using “MarkDuplicates” from Picard-tools version 1.95 (http://broadinstitute.github.io/picard/). The quality of aligned reads was examined using Ataqv v.1.0.0 ^51^. After preprocessing and quality filtering, peaks were called on alignments with MACS version 2.1.0.20151222^52^ using the parameters “-p 0.0001 -g mm -f BAMPE -- nomodel --nolambda -B --keep-dup all --call-summits”. Peaks located in blacklisted regions of the genome were removed. Overlapping peaks from all samples were merged with BEDTools version 2.26.0 53 resulting in 60,449 consensus peak regions. Raw read counts in these peaks for each sample were determined using the R package DiffBind_2.4.8 ^54^. To identify peaks with differing chromatin accessibility between WT and KO samples, the log mean normalized reads (log2 counts per million) were compared between the two sample groups. Peaks with a differing accessibility > 1.5 fold in either group were used for transcription factor motif enrichment analysis. The HOMER suite version 4.6 ^55^ and “findMotifsGenome.pl” script with parameters “mm10 -size 200” was used to determine transcription factor motifs enriched (q-value < 0.05) in the peaks of interest.

### Data Availability

The accession numbers for ATAC-seq and RNA-seq data reported in this paper are: NCBI BioProject accession PRJNA630597

### Statistical Analysis

Unless otherwise indicated, data are expressed as means ± SEM. Differences between experimental groups and time points were compared using non-parametric tests (Mann-Whitney U test), one-way or two-way ANOVA, or Student’s t test with an FDR of 5%. Statistical analyses were done with GraphPad Prism 8.4.0. Differences with a p value < 0.05 considered statistically significant.

## Acknowledgements

We thank Lesley Devine, Chao Wang for cell sorting. We thank Mei Zhong from Yale Stem Cell Center Genomics Core facility for RNA-seq library preparation and sequencing. This work was supported by Grants DK057846 and DK045735 from the NIH and SRA2014-142-S-B from the Juvenile Diabetes Research Foundation.

## Author Contributions

J.R. and K.C.H. conceived and designed the experiments and analyzed data. J.R., and S.D. conducted most of the cellular and animal experiments. G.P. and M.L.-R. labeled and viewed the human pancreas slides with the supervision from D.P. R. K. and N. L. processed the samples for ATAC-seq and analyzed the data with the supervision from M.L.S. T.S. prepared RNA and cDNA amplification for RNA-seq libraries. A.L.P. acquired the human pancreatitis slides and prepared the MHCI tetramer for IGRP staining. M.L.S., D.P. and J.L. provided valuable inputs. J.R. and K.C.H. wrote the paper with input from all the authors.

## Supplementary Information

<u>Supplementary Table 1:</u> Homer motif analysis of the transcription factors that potentially bind the chromatin sites with reduced accessibility in β cells from KO recipients.

<u>Supplementary Table 2:</u> Homer motif analysis of the transcription factors that potentially bind the chromatin sites with increased accessibility in β cells from KO recipients.

<u>Supplemental Table 3:</u> Primer pairs used for qRT-PCR analysis

<u>Supplementary Fig. 1, related to Fig 3: Normal glucose tolerance and altered islets composition in Tet2 deficient mice</u> (a) AUC glucose level in Tet2-WT and KO mice (B6 background) was measured every two weeks during IPGTTs from 4 weeks to 16 weeks of age (Data are mean ± SEM; n=6-8 mice each group). (b) H&E staining of the pancreas slides showing one islet from Tet2-WT vs KO B6 mice. Data represent at least 3 mice from each group. (c) β as well as α cells compartment in islets show as percentage of total islet cells, following intracellular staining with antibodies against insulin and glucagon and FACS analysis. Data are from ≥ 5 experiments. Each circle represents a mouse. (d) Median fluorescence intensity (MFI) showing Insulin, Nkx6.1 as well as Pdx1 content in β cells from Tet2-WT and KO mice following intracellular staining and FACS analysis. Data are from ≥ 5 experiments. One circle represents one mouse. (e) Transcriptional analysis of genes that are critical for β cell function and identity in β cells from KO vs WT mice. Insulin+ β cells were sorted following intracellular staining and mRNA level was determined after RNA recovery and cDNA synthesis. Data are from 4 sortings, n=4-5 mice each group per sorting. (*p<0.05, ***p<0.001, ****p<0.0001, ANOVA).

<u>Supplementary Fig. 2, related to Fig 3: Diabetes incidence in Tet2-WT and KO NOD mice.</u> Tet2-WT, HET and KO NOD mice were followed twice weekly for diabetes incidence up to 40 weeks of age. Results are from 11 WT, 14 HET and 17 KO female NOD mice (Long-rank curve comparison, p<0.0001).

<u>Supplementary Fig. 3, related to Fig.3: β cells from Tet2-KO mice are not protected from streptozocin killing.</u> Diabetes incidence in Tet2-WT and KO B6 mice following streptozocin (STZ) treatment. Mice were given a single dose of STZ (200 mg/kg, i.p) and followed daily for diabetes. Data are from 2 experiments and 3 mice per group per experiment (Long-rank curve comparison, p=ns).

<u>Supplementary Fig. 4, related to Fig. 4: *Tet2* gene transcription analysis in the novel subpopulation of β cells (Btm) vs normal β cells (Top).</u> Top and Btm β cells were sorted from 10-wk-old NOD mice based on Zinc and TMRE staining. RNA was recovered and the *Tet2* transcription level was measured by qRT-PCR. Results are mean ± SEM of 3 sortings, *n*=3 mice each sorting (\*\**p*<0.001, ANOVA).

<u>Supplementary Fig. 5, related to Fig. 5: The level of CD45+ infiltrates in islets from KO vs WT BMT recipients from WT NOD mice.</u> 8 weeks post-BMT, CD45+ % of total cells in the islets was analyzed by flow and collected for transcriptional profiling by Nanostring while β cells (Zinc+TMRE+) were enriched for RNA-seq as well as ATAC-seq analysis. One dot represents one recipient (Data are mean ± SEM; t-test, p=ns)

<u>Supplementary Fig. 6, related to Fig. 6:</u> Heat map generated from the 333 genes that are differently expressed between β cells from KO vs WT BMT recipients from WT NOD by bulk RNA-seq analysis (FDR<0.05).

<u>Supplementary Fig. 7, related to Fig. 6: The level of MHCI components on β cells from Tet2-KO vs WT B6 mice.</u> Histogram showing the surface level of MHCI components on β cells from B6 WT and KO mice analyzed by FACS. WT lymphocytes were included as positive control vs Isotype negative control. Data represent 3 experiments and a pair of KO and WT mice each experiment.

<u>Supplementary Fig. 8, related to Fig. 7: In vitro cytokine response from Tet2-KO vs WT islets.</u> (a) The inducible cytokines and chemokines in islets following cytokine culture. Islets from B6 mice were cultured in the presence of either IFNγ or TNFα+IL-1β+IFNγ for 24 hrs and the transcription profile of candidate chemokine and cytokine species was determined by qRT-PCR. Data are mean ± SEM of 4 independent experiment, 4 mice per experiment. (b, c) Islets from KO vs WT NOD mice of 4-5 weeks of age (b) or 6-8-week-old B6 mice (c) were cultured in the presence of IFNγ titrated at different concentrations as indicated for 24 hrs and the induction of IFNγ responsive genes as shown was analyzed and compared between KO and WT samples. Data are mean ± SEM of 3 independent experiment, 6 WT and KO mice were used each experiment. (**p<0.0001, ***p<0.001, ANOVA).

