## Supplementary Table 1 for "Tet2 Controls β cells Responses to Inflammation in Type 1 Diabetes"

| Supplementary Table 1. Homer motif analysis of the transcription factors that potentially bind the chromatin sites with reduced accessibility in β cells from KO recipients. | | | | | |
|---|---|---|---|---|---|
| Rank | Motif Name | Consensus | P-value | Log P-value | q-value (Benjamini) |
| 1 | IRF2(IRF)/Erythroblas-IRF2-ChIP-Seq(GSE36985)/Homer | GAAASYGAAASY | 1.00E-17 | -3.92E+01 | 0 |
| 2 | Foxa3(Forkhead)/Liver-Foxa3-ChIP-Seq(GSE77670)/Homer | BSNTGTTTACWYWGN | 1.00E-16 | -3.87E+01 | 0 |
| 3 | Foxa2(Forkhead)/Liver-Foxa2-ChIP-Seq(GSE25694)/Homer | CYTGTTTACWYW | 1.00E-14 | -3.31E+01 | 0 |
| 4 | IRF1(IRF)/PBMC-IRF1-ChIP-Seq(GSE43036)/Homer | GAAAGTGAAAGT | 1.00E-13 | -3.09E+01 | 0 |
| 5 | Nkx6.1(Homeobox)/Islet-Nkx6.1-ChIP-Seq(GSE40975)/Homer | GKTAATGR | 1.00E-12 | -2.77E+01 | 0 |
| 6 | Fox:Ebox(Forkhead,bHLH)/Panc1-Foxa2-ChIP-Seq(GSE47459)/Homer | NNNVCTGWGYAAACASN | 1.00E-11 | -2.62E+01 | 0 |
| 7 | FOXM1(Forkhead)/MCF7-FOXM1-ChIP-Seq(GSE72977)/Homer | TRTTTACTTW | 1.00E-11 | -2.57E+01 | 0 |
| 8 | ISRE(IRF)/ThioMac-LPS-Expression(GSE23622)/Homer | AGTTTCASTTTC | 1.00E-11 | -2.56E+01 | 0 |
| 9 | Lhx2(Homeobox)/HFSC-Lhx2-ChIP-Seq(GSE48068)/Homer | TAATTAGN | 1.00E-10 | -2.49E+01 | 0 |
| 10 | FOXA1(Forkhead)/MCF7-FOXA1-ChIP-Seq(GSE26831)/Homer | WAAGTAAACA | 1.00E-10 | -2.41E+01 | 0 |
| 11 | FOXA1(Forkhead)/LNCAP-FOXA1-ChIP-Seq(GSE27824)/Homer | WAAGTAAACA | 1.00E-09 | -2.28E+01 | 0 |
| 12 | Lhx3(Homeobox)/Neuron-Lhx3-ChIP-Seq(GSE31456)/Homer | ADBTAATTAR | 1.00E-09 | -2.25E+01 | 0 |
| 13 | Lhx1(Homeobox)/EmbryoCarcinoma-Lhx1-ChIP-Seq(GSE70957)/Homer | NNYTAATTAR | 1.00E-09 | -2.13E+01 | 0 |
| 14 | Foxo1(Forkhead)/RAW-Foxo1-ChIP-Seq(Fan_et_al.)/Homer | CTGTTTAC | 1.00E-08 | -2.00E+01 | 0 |
| 15 | IRF8(IRF)/BMDM-IRF8-ChIP-Seq(GSE77884)/Homer | GRAASTGAAAST | 1.00E-08 | -1.93E+01 | 0 |
| 16 | IRF3(IRF)/BMDM-Irf3-ChIP-Seq(GSE67343)/Homer | AGTTTCAKTTTC | 1.00E-08 | -1.88E+01 | 0 |
| 17 | FOXK2(Forkhead)/U2OS-FOXK2-ChIP-Seq(E-MTAB-2204)/Homer | SCHTGTTTACAT | 1.00E-07 | -1.75E+01 | 0 |
| 18 | Isl1(Homeobox)/Neuron-Isl1-ChIP-Seq(GSE31456)/Homer | CTAATKGV | 1.00E-07 | -1.69E+01 | 0 |
| 19 | FOXK1(Forkhead)/HEK293-FOXK1-ChIP-Seq(GSE51673)/Homer | NVWTGTTTAC | 1.00E-07 | -1.69E+01 | 0 |
| 20 | Foxo3(Forkhead)/U2OS-Foxo3-ChIP-Seq(E-MTAB-2701)/Homer | DGTAAACA | 1.00E-07 | -1.67E+01 | 0 |
| 21 | NeuroG2(bHLH)/Fibroblast-NeuroG2-ChIP-Seq(GSE75910)/Homer | ACCATCTGTT | 1.00E-06 | -1.58E+01 | 0 |
| 22 | Fra2(bZIP)/Striatum-Fra2-ChIP-Seq(GSE43429)/Homer | GGATGACTCATC | 1.00E-06 | -1.57E+01 | 0 |
| 23 | NeuroD1(bHLH)/Islet-NeuroD1-ChIP-Seq(GSE30298)/Homer | GCCATCTGTT | 1.00E-06 | -1.56E+01 | 0 |
| 24 | Stat3(Stat)/mES-Stat3-ChIP-Seq(GSE11431)/Homer | CTTCCGGGAA | 1.00E-06 | -1.55E+01 | 0 |
| 25 | Fra1(bZIP)/BT549-Fra1-ChIP-Seq(GSE46166)/Homer | NNATGASTCATH | 1.00E-06 | -1.52E+01 | 0 |
| 26 | Six1(Homeobox)/Myoblast-Six1-ChIP-Chip(GSE20150)/Homer | GKVTCADRTTWC | 1.00E-06 | -1.48E+01 | 0 |
| 27 | Nanog(Homeobox)/mES-Nanog-ChIP-Seq(GSE11724)/Homer | RGCCATTAAC | 1.00E-06 | -1.48E+01 | 0 |
| 28 | SCL(bHLH)/HPC7-Scl-ChIP-Seq(GSE13511)/Homer | AVCAGCTG | 1.00E-06 | -1.47E+01 | 0 |
| 29 | Tcf21(bHLH)/ArterySmoothMuscle-Tcf21-ChIP-Seq(GSE61369)/Homer | NAACAGCTGG | 1.00E-06 | -1.43E+01 | 0 |
| 30 | Atoh1(bHLH)/Cerebellum-Atoh1-ChIP-Seq(GSE22111)/Homer | VNRVCAGCTGGY | 1.00E-05 | -1.37E+01 | 0 |
| 31 | Pdx1(Homeobox)/Islet-Pdx1-ChIP-Seq(SRA008281)/Homer | YCATYAATCA | 1.00E-05 | -1.36E+01 | 0 |
| 32 | AP-1(bZIP)/ThioMac-PU.1-ChIP-Seq(GSE21512)/Homer | VTGACTCATC | 1.00E-05 | -1.36E+01 | 0 |
| 33 | FoxL2(Forkhead)/Ovary-FoxL2-ChIP-Seq(GSE60858)/Homer | WWTRTAAACAVG | 1.00E-05 | -1.36E+01 | 0 |
| 34 | Stat3+il21(Stat)/CD4-Stat3-ChIP-Seq(GSE19198)/Homer | SVYTTCCNGGAARB | 1.00E-05 | -1.35E+01 | 0 |
| 35 | HEB(bHLH)/mES-Heb-ChIP-Seq(GSE53233)/Homer | VCAGCTGBNN | 1.00E-05 | -1.35E+01 | 0 |
| 36 | Rfx6(HTH)/Min6b1-Rfx6.HA-ChIP-Seq(GSE62844)/Homer | TGTTKCCTAGCAACM | 1.00E-05 | -1.32E+01 | 0 |
| 37 | MyoG(bHLH)/C2C12-MyoG-ChIP-Seq(GSE36024)/Homer | AACAGCTG | 1.00E-05 | -1.31E+01 | 0 |
| 38 | JunB(bZIP)/DendriticCells-Junb-ChIP-Seq(GSE36099)/Homer | RATGASTCAT | 1.00E-05 | -1.31E+01 | 0 |

| | | | | | |
|---|---|---|---|---|---|
| 39 | FOXP1(Forkhead)/H9-FOXP1-ChIP-Seq(GSE31006)/Homer | NYYTGTTTACHN | 1.00E-05 | -1.25E+01 | 0 |
| 40 | BATF(bZIP)/Th17-BATF-ChIP-Seq(GSE39756)/Homer | DATGASTCAT | 1.00E-05 | -1.23E+01 | 0 |
| 41 | NF1(CTF)/LNCAP-NF1-ChIP-Seq(Unpublished)/Homer | CYTGGCABNSTGCCAR | 1.00E-05 | -1.23E+01 | 0 |
| 42 | Foxf1(Forkhead)/Lung-Foxf1-ChIP-Seq(GSE77951)/Homer | WWATRTAAACAN | 1.00E-05 | -1.22E+01 | 0 |
| 43 | Ap4(bHLH)/AML-Tfap4-ChIP-Seq(GSE45738)/Homer | NAHCAGCTGD | 1.00E-05 | -1.18E+01 | 0.0001 |
| 44 | Ptf1a(bHLH)/Panc1-Ptf1a-ChIP-Seq(GSE47459)/Homer | ACAGCTGTTN | 1.00E-05 | -1.17E+01 | 0.0001 |
| 45 | Rfx5(HTH)/GM12878-Rfx5-ChIP-Seq(GSE31477)/Homer | SCCTAGCAACAG | 1.00E-05 | -1.17E+01 | 0.0001 |
| 46 | Atf3(bZIP)/GBM-ATF3-ChIP-Seq(GSE33912)/Homer | DATGASTCATHN | 1.00E-05 | -1.17E+01 | 0.0001 |
| 47 | Ascl1(bHLH)/NeuralTubes-Ascl1-ChIP-Seq(GSE55840)/Homer | NNVVCAGCTGBN | 1.00E-04 | -1.13E+01 | 0.0001 |
| 48 | NFkB-p65(RHD)/GM12787-p65-ChIP-Seq(GSE19485)/Homer | WGGGGATTTCCC | 1.00E-04 | -1.05E+01 | 0.0002 |
| 49 | Atf7(bZIP)/3T3L1-Atf7-ChIP-Seq(GSE56872)/Homer | NGRTGACGTCAY | 1.00E-04 | -1.05E+01 | 0.0002 |
| 50 | STAT1(Stat)/HelaS3-STAT1-ChIP-Seq(GSE12782)/Homer | NATTTCCNGGAAAT | 1.00E-04 | -1.02E+01 | 0.0003 |
| 51 | Mef2d(MADS)/Retina-Mef2d-ChIP-Seq(GSE61391)/Homer | GCTATTTTTAGC | 1.00E-04 | -1.01E+01 | 0.0003 |
| 52 | E2A(bHLH)/proBcell-E2A-ChIP-Seq(GSE21978)/Homer | DNRCAGCTGY | 1.00E-04 | -9.97E+00 | 0.0003 |
| 53 | IRF4(IRF)/GM12878-IRF4-ChIP-Seq(GSE32465)/Homer | ACTGAAACCA | 1.00E-04 | -9.92E+00 | 0.0003 |
| 54 | Fosl2(bZIP)/3T3L1-Fosl2-ChIP-Seq(GSE56872)/Homer | NATGASTCABNN | 1.00E-04 | -9.70E+00 | 0.0004 |
| 55 | MafA(bZIP)/Islet-MafA-ChIP-Seq(GSE30298)/Homer | TGCTGACTCA | 1.00E-04 | -9.40E+00 | 0.0005 |
| 56 | PRDM1(Zf)/Hela-PRDM1-ChIP-Seq(GSE31477)/Homer | ACTTTCACTTTC | 1.00E-03 | -9.11E+00 | 0.0007 |
| 57 | NFkB-p65-Rel(RHD)/ThioMac-LPS-Expression(GSE23622)/Homer | GGAAATTCCC | 1.00E-03 | -8.98E+00 | 0.0008 |
| 58 | STAT4(Stat)/CD4-Stat4-ChIP-Seq(GSE22104)/Homer | NYTTCCWGGAAR | 1.00E-03 | -8.89E+00 | 0.0009 |
| 59 | BMYB(HTH)/Hela-BMYB-ChIP-Seq(GSE27030)/Homer | NHAACBGYYV | 1.00E-03 | -8.60E+00 | 0.0011 |
| 60 | Tbr1(T-box)/Cortex-Tbr1-ChIP-Seq(GSE71384)/Homer | AAGGTGTKAA | 1.00E-03 | -8.44E+00 | 0.0013 |
| 61 | NF1-halfsite(CTF)/LNCaP-NF1-ChIP-Seq(Unpublished)/Homer | YTGCCAAG | 1.00E-03 | -8.28E+00 | 0.0015 |
| 62 | Atf2(bZIP)/3T3L1-Atf2-ChIP-Seq(GSE56872)/Homer | NRRTGACGTCAT | 1.00E-03 | -8.23E+00 | 0.0016 |
| 63 | PU.1:IRF8(ETS:IRF)/pDC-Irf8-ChIP-Seq(GSE66899)/Homer | GGAAGTGAAAST | 1.00E-03 | -7.43E+00 | 0.0034 |
| 64 | Jun-AP1(bZIP)/K562-cJun-ChIP-Seq(GSE31477)/Homer | GATGASTCATCN | 1.00E-03 | -7.29E+00 | 0.0039 |
| 65 | MyoD(bHLH)/Myotube-MyoD-ChIP-Seq(GSE21614)/Homer | RRCAGCTGYTSY | 1.00E-03 | -7.14E+00 | 0.0045 |
| 66 | Tcf12(bHLH)/GM12878-Tcf12-ChIP-Seq(GSE32465)/Homer | VCAGCTGYTG | 1.00E-03 | -7.12E+00 | 0.0045 |
| 67 | MITF(bHLH)/MastCells-MITF-ChIP-Seq(GSE48085)/Homer | RTCATGTGAC | 1.00E-03 | -7.02E+00 | 0.0049 |
| 68 | Unknown-ESC-element(?)/mES-Nanog-ChIP-Seq(GSE11724)/Homer | CACAGCAGGGGG | 1.00E-02 | -6.86E+00 | 0.0056 |
| 69 | c-Jun-CRE(bZIP)/K562-cJun-ChIP-Seq(GSE31477)/Homer | ATGACGTCATCY | 1.00E-02 | -6.74E+00 | 0.0063 |
| 70 | Hoxb4(Homeobox)/ES-Hoxb4-ChIP-Seq(GSE34014)/Homer | TGATTRATGGCY | 1.00E-02 | -6.69E+00 | 0.0064 |
| 71 | Bach2(bZIP)/OCILy7-Bach2-ChIP-Seq(GSE44420)/Homer | TGCTGAGTCA | 1.00E-02 | -6.49E+00 | 0.0078 |
| 72 | Atf1(bZIP)/K562-ATF1-ChIP-Seq(GSE31477)/Homer | GATGACGTCA | 1.00E-02 | -6.41E+00 | 0.0083 |
| 73 | CRE(bZIP)/Promoter/Homer | CSGTGACGTCAC | 1.00E-02 | -6.40E+00 | 0.0083 |
| 74 | USF1(bHLH)/GM12878-Usf1-ChIP-Seq(GSE32465)/Homer | SGTCACGTGR | 1.00E-02 | -6.39E+00 | 0.0083 |
| 75 | Nur77(NR)/K562-NR4A1-ChIP-Seq(GSE31363)/Homer | TGACCTTTNCNT | 1.00E-02 | -6.19E+00 | 0.01 |
| 76 | Olig2(bHLH)/Neuron-Olig2-ChIP-Seq(GSE30882)/Homer | RCCATMTGTT | 1.00E-02 | -6.17E+00 | 0.0101 |
| 77 | Mef2b(MADS)/HEK293-Mef2b.V5-ChIP-Seq(GSE67450)/Homer | GCTATTTTTGGM | 1.00E-02 | -6.13E+00 | 0.0103 |
| 78 | Usf2(bHLH)/C2C12-Usf2-ChIP-Seq(GSE36030)/Homer | GTCACGTGGT | 1.00E-02 | -5.71E+00 | 0.0155 |
| 79 | HNF1b(Homeobox)/PDAC-HNF1B-ChIP-Seq(GSE64557)/Homer | GTTAATNATTAA | 1.00E-02 | -5.56E+00 | 0.0177 |

| | | | | | |
|---|---|---|---|---|---|
| 80 | Tbet(T-box)/CD8-Tbet-ChIP-Seq(GSE33802)/Homer | AGGTGTGAAM | 1.00E-02 | -5.55E+00 | 0.0177 |
| 81 | Tgif2(Homeobox)/mES-Tgif2-ChIP-Seq(GSE55404)/Homer | TGTCANYT | 1.00E-02 | -5.54E+00 | 0.0177 |
| 82 | Tbx5(T-box)/HL1-Tbx5.biotin-ChIP-Seq(GSE21529)/Homer | AGGTGTCA | 1.00E-02 | -5.45E+00 | 0.019 |
| 83 | STAT5(Stat)/mCD4+-Stat5-ChIP-Seq(GSE12346)/Homer | RTTTCTNAGAAA | 1.00E-02 | -5.41E+00 | 0.0196 |
| 84 | Myf5(bHLH)/GM-Myf5-ChIP-Seq(GSE24852)/Homer | BAACAGCTGT | 1.00E-02 | -5.40E+00 | 0.0196 |
| 85 | Fli1(ETS)/CD8-FLI-ChIP-Seq(GSE20898)/Homer | NRYTTCCGGH | 1.00E-02 | -5.35E+00 | 0.0203 |
| 86 | Six2(Homeobox)/NephronProgenitor-Six2-ChIP-Seq(GSE39837)/Homer | GWAAYHTGAKMC | 1.00E-02 | -5.33E+00 | 0.0205 |
| 87 | Esrrb(NR)/mES-Esrrb-ChIP-Seq(GSE11431)/Homer | KTGACCTTGA | 1.00E-02 | -5.00E+00 | 0.0282 |
| 88 | Zic3(Zf)/mES-Zic3-ChIP-Seq(GSE37889)/Homer | GGCCYCCTGCTGDGH | 1.00E-02 | -5.00E+00 | 0.0282 |
| 89 | BORIS(Zf)/K562-CTCFL-ChIP-Seq(GSE32465)/Homer | CNNBRGCGCCCCCTGSTGGC | 1.00E-02 | -4.91E+00 | 0.0301 |
| 90 | Mef2a(MADS)/HL1-Mef2a.biotin-ChIP-Seq(GSE21529)/Homer | CYAAAAATAG | 1.00E-02 | -4.88E+00 | 0.0306 |
| 91 | Srebp1a(bHLH)/HepG2-Srebp1a-ChIP-Seq(GSE31477)/Homer | RTCACSCCAY | 1.00E-02 | -4.80E+00 | 0.033 |
| 92 | c-Myc(bHLH)/LNCAP-cMyc-ChIP-Seq(Unpublished)/Homer | VCCACGTG | 1.00E-02 | -4.79E+00 | 0.033 |
| 93 | HOXA2(Homeobox)/mES-Hoxa2-ChIP-Seq(Donaldson_et_al.)/Homer | GYCATCMATCAT | 1.00E-02 | -4.76E+00 | 0.0337 |
| 94 | Elk4(ETS)/Hela-Elk4-ChIP-Seq(GSE31477)/Homer | NRYTTCCGGY | 1.00E-01 | -4.57E+00 | 0.0402 |
| 95 | CLOCK(bHLH)/Liver-Clock-ChIP-Seq(GSE39860)/Homer | GHCACGTG | 1.00E-01 | -4.49E+00 | 0.0432 |
| 96 | Zic(Zf)/Cerebellum-ZIC1.2-ChIP-Seq(GSE60731)/Homer | CCTGCTGAGH | 1.00E-01 | -4.38E+00 | 0.0475 |
| 97 | IRF:BATF(IRF:bZIP)/pDC-Irf8-ChIP-Seq(GSE66899)/Homer | CTTTCANTATGACTV | 1.00E-01 | -4.35E+00 | 0.0483 |
