## Supplementary Table 2 for "Tet2 Controls β cells Responses to Inflammation in Type 1 Diabetes"

| Rank | Motif Name | Consensus | P-value | Log P-value | q-value (Benjamini) |
|---|---|---|---|---|---|
| 1 | CTCF(Zf)/CD4+-CTCF-ChIP-Seq(Barski_et_al.)/Homer | AYAGTGCCMYCTRGTGGCCA | 1.00E-05 | -1.33E+01 | 0.0006 |
| 2 | Fox:Ebox(Forkhead,bHLH)/Panc1-Foxa2-ChIP-Seq(GSE47459)/Homer | NNNVCTGWGYAAACASN | 1.00E-05 | -1.15E+01 | 0.0018 |
| 3 | FOXA1(Forkhead)/MCF7-FOXA1-ChIP-Seq(GSE26831)/Homer | WAAGTAAACA | 1.00E-04 | -1.01E+01 | 0.0052 |
| 4 | Foxa3(Forkhead)/Liver-Foxa3-ChIP-Seq(GSE77670)/Homer | BSNTGTTTACWYWGN | 1.00E-04 | -9.92E+00 | 0.0052 |
| 5 | FOXM1(Forkhead)/MCF7-FOXM1-ChIP-Seq(GSE72977)/Homer | TRTTTACTTW | 1.00E-04 | -9.52E+00 | 0.0054 |
| 6 | FOXA1(Forkhead)/LNCAP-FOXA1-ChIP-Seq(GSE27824)/Homer | WAAGTAAACA | 1.00E-03 | -9.08E+00 | 0.0069 |
| 7 | Pdx1(Homeobox)/Islet-Pdx1-ChIP-Seq(SRA008281)/Homer | YCATYAATCA | 1.00E-03 | -8.34E+00 | 0.0125 |
| 8 | BORIS(Zf)/K562-CTCFL-ChIP-Seq(GSE32465)/Homer | CNNBRGCGCCCCCTGSTGGC | 1.00E-03 | -8.27E+00 | 0.0125 |
| 9 | Foxo3(Forkhead)/U2OS-Foxo3-ChIP-Seq(E-MTAB-2701)/Homer | DGTAAACA | 1.00E-03 | -8.25E+00 | 0.0125 |
| 10 | X-box(HTH)/NPC-H3K4me1-ChIP-Seq(GSE16256)/Homer | GGTTGCCATGGCAA | 1.00E-03 | -8.06E+00 | 0.0125 |
| 11 | Foxa2(Forkhead)/Liver-Foxa2-ChIP-Seq(GSE25694)/Homer | CYTGTTTACWYW | 1.00E-03 | -7.82E+00 | 0.0133 |
| 12 | Oct6(POU,Homeobox)/NPC-Pou3f1-ChIP-Seq(GSE35496)/Homer | WATGCAAATGAG | 1.00E-03 | -7.53E+00 | 0.0163 |
| 13 | Brn1(POU,Homeobox)/NPC-Brn1-ChIP-Seq(GSE35496)/Homer | TATGCWAATBAV | 1.00E-03 | -7.11E+00 | 0.0229 |
| 14 | Rfx1(HTH)/NPC-H3K4me1-ChIP-Seq(GSE16256)/Homer | KGTTGCCATGGCAA | 1.00E-02 | -6.27E+00 | 0.0492 |
| 15 | Lhx3(Homeobox)/Neuron-Lhx3-ChIP-Seq(GSE31456)/Homer | ADBTAATTAR | 1.00E-02 | -6.26E+00 | 0.0492 |

Table S2. Homer motif analysis of the transcription factors that potentially bind the chromatin sites with increased accessibility in β cells from KO recipients.
