## Supplementary Table 3 for "Tet2 Controls β cells Responses to Inflammation in Type 1 Diabetes"

**Supplementary Table 3: Primer pairs used for qRT-PCR a**

| | |
|---|---|
| *TET2-F* | GGCTACAAAGCTCCAGAATGG |
| *TET2-R* | AAGAGTGCCACTTGGTGTCTC |
| *Tet2-F* | AGAGAAGACAATCGAGAAGTCGG |
| *Tet2-R* | CCTTCCGTACTCCCAAACTCAT |
| *Il6-F* | TCTATACCACTTCACAAGTCGGA |
| *Il6-R* | GAATTGCCATTGCACAACTCTTT |
| *Ccl2-F* | TTAAAAACCTGGATCGGAACCAA |
| *Ccl2-R* | GCATTAGCTTCAGATTTACGGGT |
| *Cxcl10-F* | CCAAGTGCTGCCGTCATTTTC |
| *Cxcl10-R* | GGCTCGCAGGGATGATTTCAA |
| *Cxcl16-F* | CCTTGTCTCTTGCGTTCTTCC |
| *Cxcl16-R* | TCCAAAGTACCCTGCGGTATC |
| *Tnf-F* | CAGGCGGTGCCTATGTCTC |
| *Tnf-R* | CGATCACCCCGAAGTTCAGTAG |
| *Fas-F* | GCGGGTTCGTGAAACTGATAA |
| *Fas-R* | GCAAAATGGGCCTCCTTGATA |
| *Tnfrsf10b-F* | CGGGCAGATCACTACACCC |
| *Tnfrsf10b-R* | TGTTACTGGAACAAAGACAGCC |
| *stat1-F* | TCACAGTGGTTCGAGCTTCAG |
| *stat1-R* | CGAGACATCATAGGCAGCGTG |
| *Ccl5-F* | GCTGCTTTGCCTACCTCTCC |
| *Ccl5-R* | TCGAGTGACAAACACGACTGC |
| *Cxcl12-F* | TGCATCAGTGACGGTAAACCA |
| *Cxcl12-R* | TTCTTCAGCCGTGCAACAATC |
| *Ccl19-F* | CCTGGGAACATCGTGAAAGC |
| *Ccl19-R* | TAGTGTGGTGAACACAACAGC |
| *Ccl20-F* | ACTGTTGCCTCTCGTACATACA |
| *Ccl20-R* | GAGGAGGTTCACAGCCCTTTT |
| *Il1b-F* | GAAATGCCACCTTTTGACAGTG |
| *Il1b-R* | TGGATGCTCTCATCAGGACAG |
| *Cxcl9-F* | GGAGTTCGAGGAACCCTAGTG |
| *Cxcl9-R* | GGGATTTGTAGTGGATCGTGC |
| *Cxcl11-F* | GGCTTCCTTATGTTCAAACAGGG |
| *Cxcl11-R* | GCCGTTACTCGGGTAAATTACA |
| *Pdl1* -F | AGTATGGCAGCAACGTCACG |
| *Pdl1* -R | TCCTTTTCCCAGTACACCACTA |
| *Ido-F* | TGGCGTATGTGTGGAACCG |
| *Ido-R* | CTCGCAGTAGGGAACAGCAA |
| *Actb-F* | GGCTGTATTCCCCTCCATCG |
| *Actb-R* | CCAGTTGGTAACAATGCCATGT |

**ınalysis**
