## Supplementary Figures for "Tet2 Controls β cells Responses to Inflammation in Type 1 Diabetes"

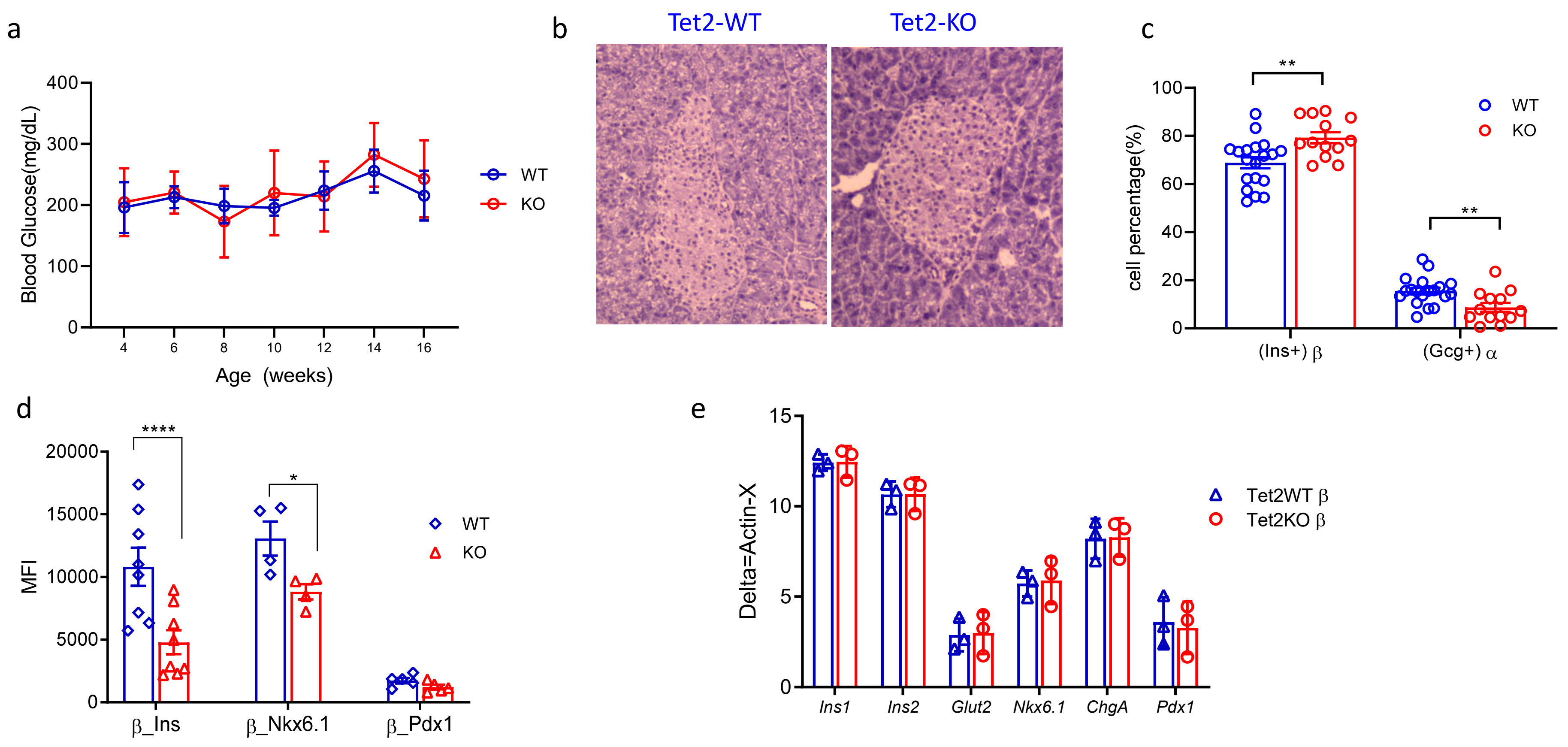


**Supplementary Fig 1, related to Fig 3. Normal glucose tolerance and altered islets composition in Tet2 deficient mice** (a) AUC glucose level in Tet2-WT and KO mice (B6 background) was measured every two weeks during IPGTTs from 4 weeks to 16 weeks of age (Data are mean ± SEM; n=6-8 mice each group). (b) H&E staining of the pancreas slides showing one islet from Tet2-WT vs KO B6 mice. Data represent at least 3 mice from each group. (c) β as well as α cells compartment in islets show as percentage of total islet cells, following intracellular staining with antibodies against insulin and glucagon and FACS analysis. Data are from ≥ 5 experiments. Each circle represents a mouse. (d) Median fluorescence intensity (MFI) showing Insulin, Nkx6.1 as well as Pdx1 content in β cells from Tet2-WT and KO mice following intracellular staining and FACS analysis. Data are from ≥ 5 experiments. One circle represents one mouse. (e) Transcriptional analysis of genes that are critical for β cell function and identity in β cells from KO vs WT mice. Insulin+ β cells were sorted following intracellular staining and mRNA level was determined after RNA recovery and cDNA synthesis. Data are from 4 sortings, n=4-5 mice each group per sorting. (*p<0.05, ***p<0.001, ****p<0.0001, ANOVA).

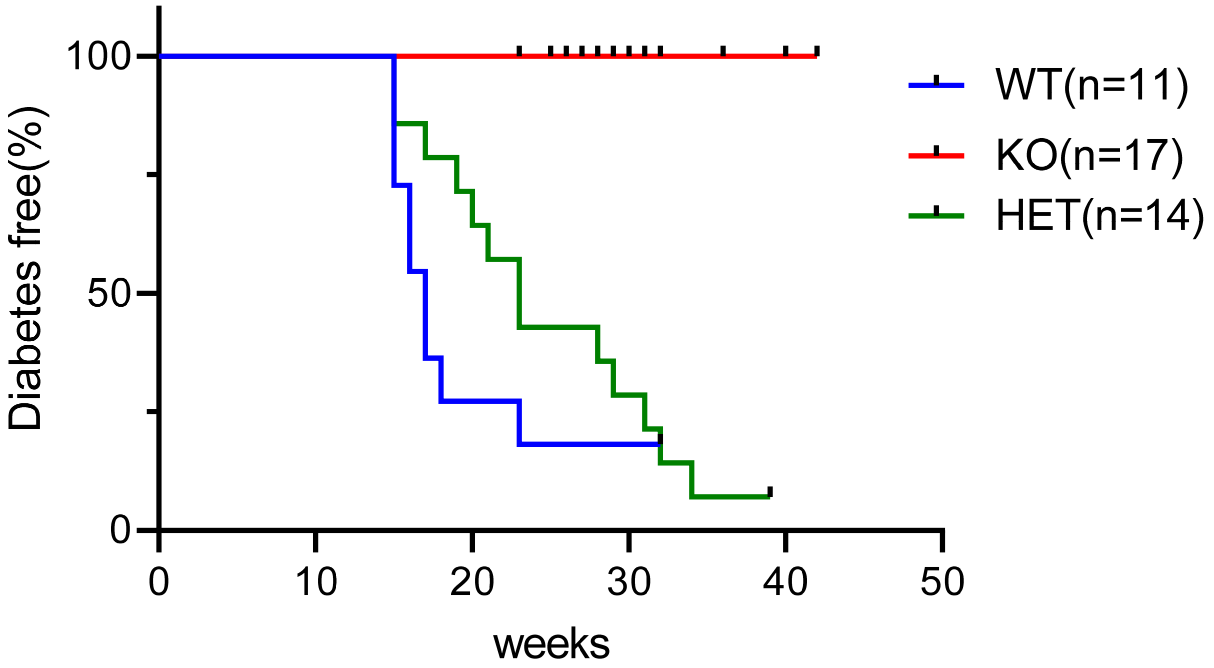


**Supplementary Fig 2, related to Fig 3**. **Diabetes incidence in Tet2-WT and KO NOD mice**. Tet2-WT , HET and KO NOD mice were followed twice weekly for diabetes incidence up to 40 weeks of age. Results are from 11 WT, 14 HET and 17 KO female NOD mice (Long-rank curve comparison, p<0.0001).

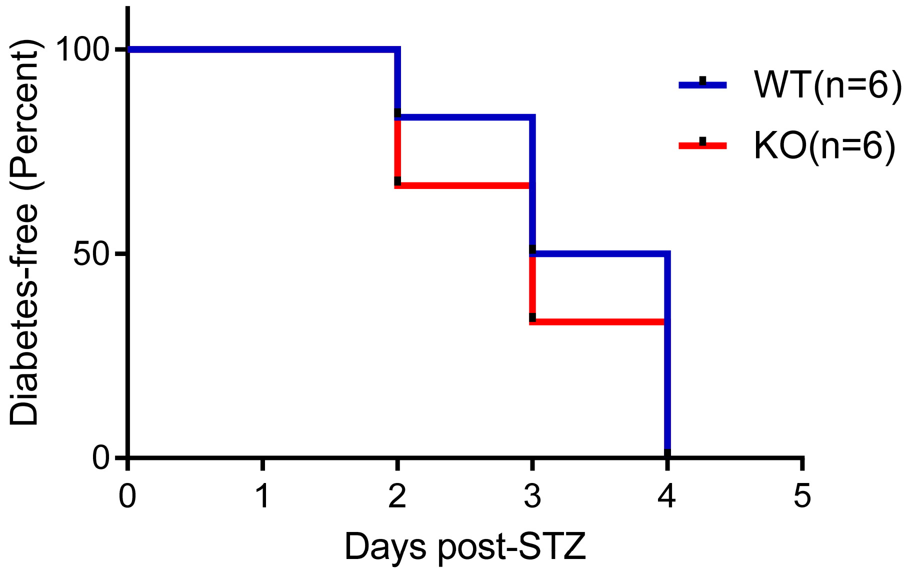


**Supplementary Fig 3, related to Fig 3**. **β cells from Tet2-KO mice are not protected from streptozocin killing.** Diabetes incidence in Tet2-WT and KO B6 mice following streptozocin (STZ) treatment. Mice were given a single dose of STZ (200 mg/kg, i.p) and followed daily for diabetes. Data are from 2 experiments and 3 mice per group per experiment (Long-rank curve comparison, p=ns).

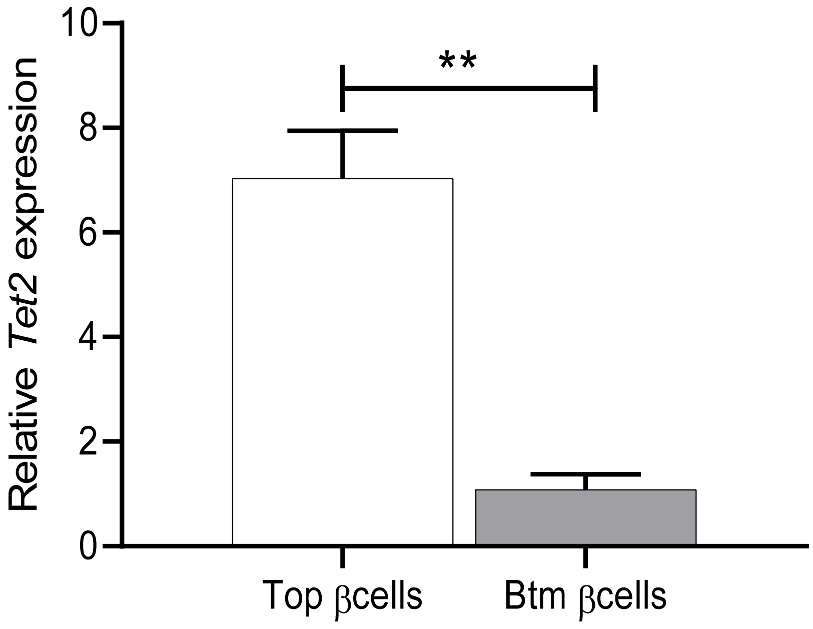


**Supplementary Fig 4, related to Fig 4**. ***Tet2* gene transcription analysis in the novel subpopulation of β cells (Btm) vs normal β cells (Top).** Top and Btm β cells were sorted from 10-wk-old NOD mice based on Zinc and TMRE staining. RNA was recovered and the *Tet2* transcription level was measured by qRT-PCR. Results are mean ± SEM of 3 sortings, $n$=3 mice each sorting (**$p$<0.001, ANOVA).

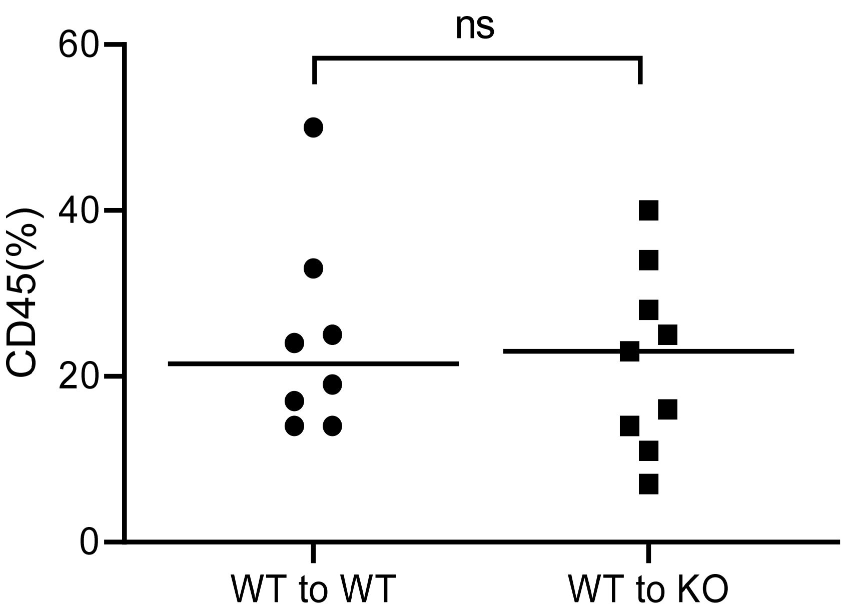


**Supplementary Fig 5, related to Fig 5**. **The level of CD45+ infiltrates in islets from KO vs WT BMT recipients from WT NOD mice**. 8 weeks post-BMT, CD45+ % of total cells in the islets was analyzed by flow and collected for transcriptional profiling by Nanostring while β cells (Zinc+TMRE+) were enriched for RNAseq as well as ATACseq analysis. One dot represents one recipient (Data are mean ± SEM; t-test, p=ns)

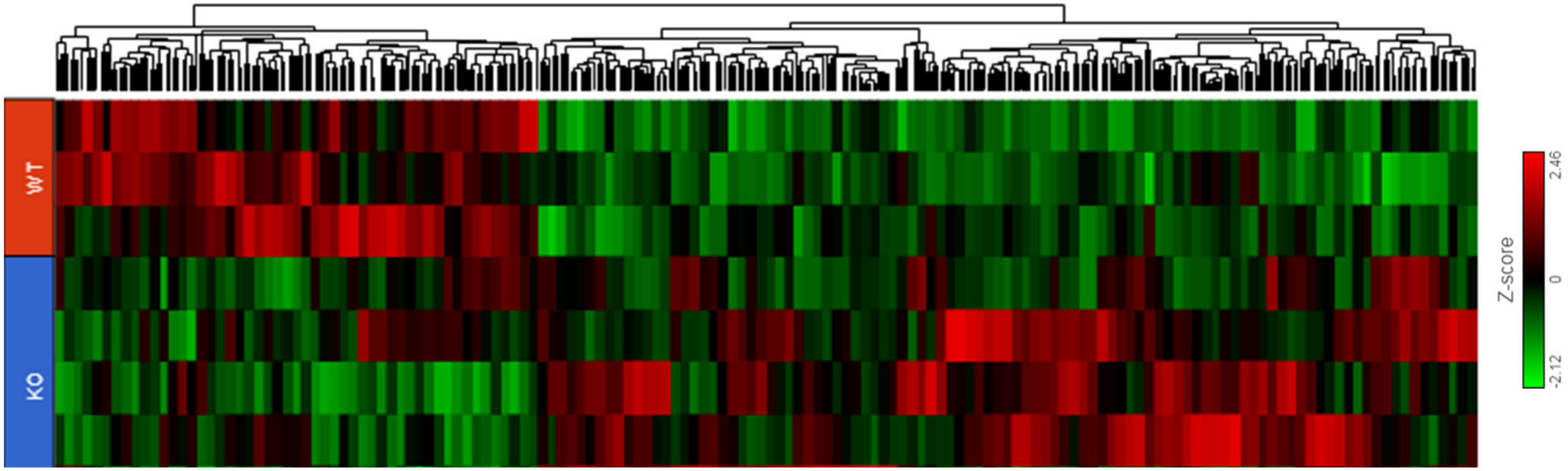


**Supplementary Fig 6, related to Fig 6**. Heat map generated from the 333 genes that are differently expressed between β cells from KO vs WT BMT recipients from WT NOD by bulk RNASEQ analysis (FDR<0.05).

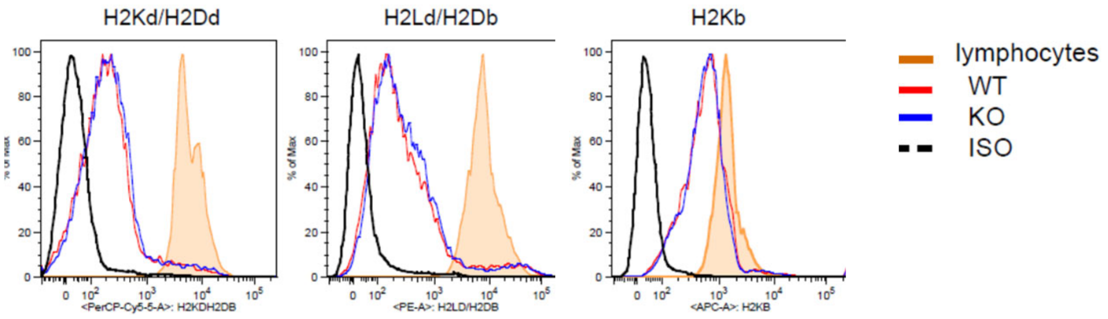


**Supplementary Fig 7, related to Fig 6**. **The level of MHCI components on β cells from Tet2-KO vs WT B6 mice**

Histogram showing the surface level of MHCI components on β cells from B6 WT and KO mice analyzed by FACS. WT lymphocytes were included as positive control vs Isotype negative control. Data represent 3 experiments and a pair of KO and WT mice each experiment.

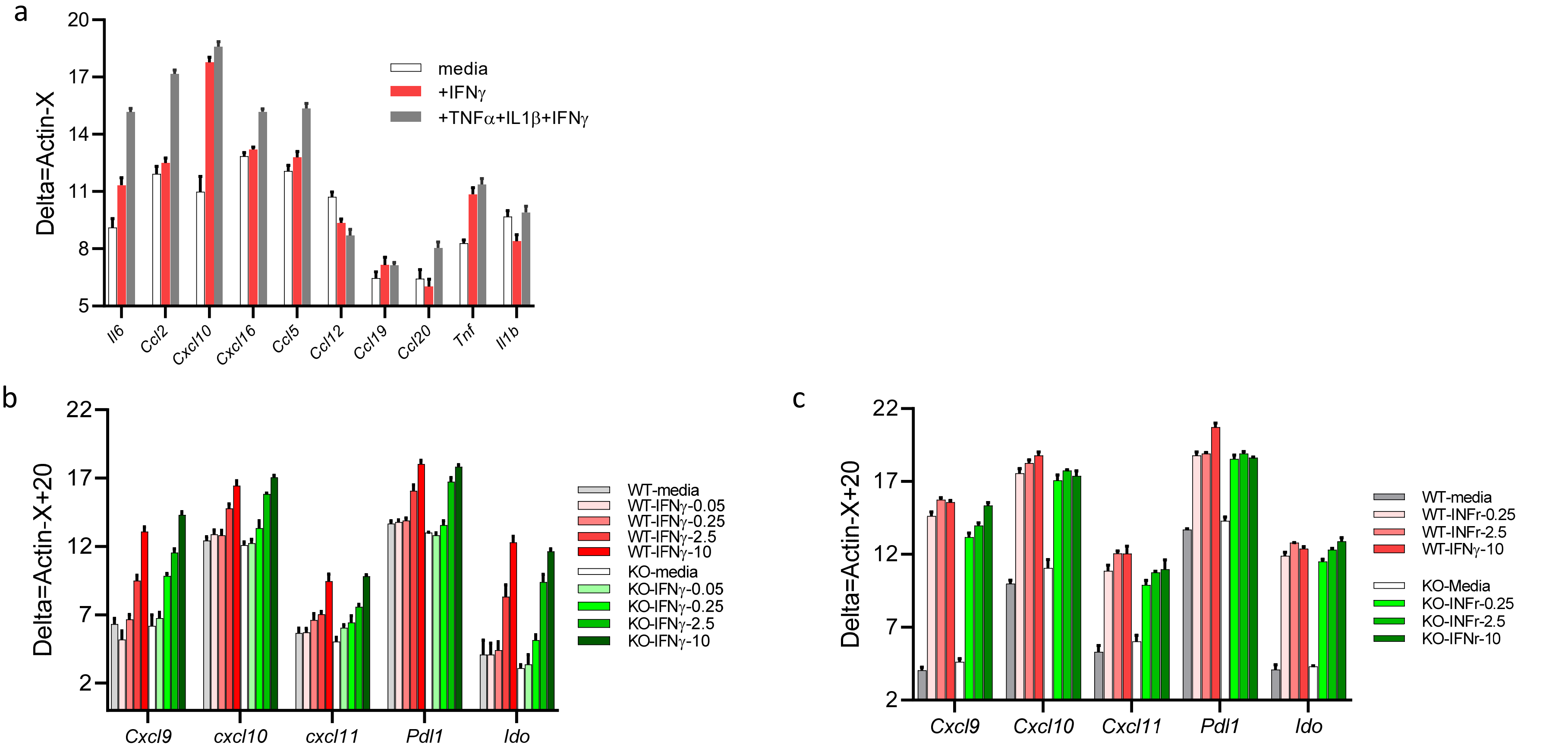


**Supplementary Fig 8, related to Fig 7**. **In vitro cytokine response from Tet2-KO vs WT islets**. (a) The inducible cytokines and chemokines in islets following cytokine culture. Islets from B6 mice were cultured in the presence of either IFNγ or TNFα+IL-1β+IFNγ for 24 hrs and the transcription profile of candidate chemokine and cytokine species was determined by qRT-PCR. Data are mean ± SEM of 4 independent experiment, 4 mice per experiment. (b and c) Islets from KO vs WT NOD mice of 4-5 weeks of age (b) or 6-8-week-old B6 mice (c) were cultured in the presence of IFNγ titrated at different concentrations as indicated for 24 hrs and the induction of IFNγ responsive genes as shown was analyzed and compared between KO and WT samples. Data are mean ± SEM of 3 independent experiment, 6 WT and KO mice were used each experiment. (**p<0.0001, ***p<0.001, ANOVA).
